# An A-rich linker between dengue virus tandem xrRNAs facilitates functional coordination

**DOI:** 10.64898/2026.03.04.709395

**Authors:** Elizabeth Spear, Zoe O’Donoghue, Steve L. Bonilla, Kevin P. Larsen, Madeline E Sherlock, Jeffrey S. Kieft

## Abstract

Orthoflaviviruses use programmed resistance to host 5′ to 3′ exoribonucleases to produce subgenomic flaviviral RNAs (sfRNAs) during infection. This resistance is conferred by exoribonuclease-resistant RNA (xrRNA) structures that often occur in tandem and whose function can be coupled. In dengue virus serotype 2 (DENV2) this coupling results in changing patterns of sfRNA identity and abundance linked to the ability of the virus to adapt to host vs. vector infections. The physical basis of this coupling was unknown. Using a combination of virology, biochemistry, bioinformatics, structural biology, and biophysics, we explored the structural and sequence determinants of tandem xrRNA coupling in DENV2. We discovered that the spatial proximity, order, and structural integrity of the tandem xrRNAs are all important for coupling. Furthermore, an unpaired A-rich linker that lies between the two xrRNAs is essential in stabilizing a specific structure that correlates to coupling. This A-rich sequence likely forms tertiary contacts with an adjacent stem-loop structure to form a physical bridge between the two xrRNAs, a finding that is supported by a mid-resolution cryoEM map of the DENV2 tandem xrRNAs. Disruption of the structure of this bridge by mutation changes the relative orientation or spacing between the tandem xrRNAs, which is correlated to their functional coupling. These findings provide an explanation for the coupling between tandem xrRNAs and suggests new mechanistic hypotheses.

**IMPORTANCE:** Dengue virus (DENV) generates non-coding subgenomic flaviviral RNAs (sfRNAs) that affect several cellular pathways and are important for successful infection. These sfRNAs are formed by structured RNA elements in the viral genome called exoribonuclease-resistant RNAs (xrRNAs), which fold into a distinct three-dimensional topology to block degradation by host cell exoribonucleases and often occur in tandem. Specific patterns of sfRNAs made during infection are important for host vs. vector fitness, and in DENV2 this pattern depends on functional coupling between tandem xrRNAs. However, the source of this functional coupling was unknown. We determined that an unpaired A-rich linker between the tandem xrRNAs is necessary for creating a structural bridge between the tandem xrRNAs. This bridge appears to favor a specific orientation between the tandem xrRNAs that is correlated to coupling and therefore to the patterns and relative abundance of sfRNAs produced during infection.

## INTRODUCTION

Mosquito-borne orthoflaviviruses (MBFV), including dengue virus (DENV), Zika virus (ZIKV), West Nile virus (WNV), Japanese encephalitis virus (JEV), yellow fever virus (YFV) and many others, are positive-sense, single-stranded RNA viruses that pose a global threat to human health (1–3). DENV alone is responsible for ∼400 million infections each year and, despite causing debilitating illness, there are limited vaccine options and no approved antiviral treatments (4). The 2015-2016 ZIKV outbreak provided a reminder that flaviviruses can rapidly emerge or re-emerge as global threats and present unexpected adverse health effects. For example, ZIKV can be sexually transmitted or vertically transmitted from mother to fetus, which then can lead to a spectrum of fetal abnormalities; similar to DENV, there are no treatments or vaccines (5). With the range of host mosquitos expanding, there is a pressing need to study and understand the basic molecular processes that underlie orthoflavivirus infection and thus inform the development of new therapies and vaccines (6).

During infection, orthoflaviviruses generate specific viral non-coding RNAs that accumulate to high levels in the cell (7, 8). These subgenomic flaviviral RNAs (sfRNAs) are several hundred nucleotides long and are derived from the 3′ untranslated region (UTR) of the viral genomic RNA (9). The sfRNAs operate as entities separate from the genome to facilitate replication, cytopathicity, and pathogenesis of orthoflaviviruses through several identified pathways (8, 10–22). These include dampening the innate immune response of both the mosquito vector and the mammalian hosts by interfering with RNAi and interferon antiviral responses, respectively (14, 23–29). In this capacity, the sfRNAs play a role in the ability of these viruses to infect and switch between both arthropod vector and vertebrate hosts and to affect cellular tropism during infection (11, 30–35). Importantly, the identity, amount, and relative abundances of different sfRNAs (the ‘sfRNA pattern’) can vary between viruses and can depend on the type of cell infected, which appears to allow the virus to adapt and maintain fitness in different hosts (32–34, 36, 37). The sfRNA-dependent functions must involve interactions with host protein factors, but the detailed molecular mechanisms are incompletely understood.

sfRNAs are formed by incomplete degradation of the viral genomic RNA by host cell 5′ to 3′ exoribonucleases, primarily Xrn1 (9). Xrn1 is the processive enzyme primarily responsible for cytoplasmic cellular RNA decay, and it has a role during innate immunity to degrade viral RNA (38, 39). During orthoflavivirus infection, Xrn1 degrades a subset of the viral genomic RNA in the cell in a 5′ to 3′ direction until it reaches specialized RNA structures located within the genome’s 3′ UTR (9, 40–42). These structures, dubbed exoribonuclease-resistant RNAs (xrRNAs), efficiently block further progression of the exoribonuclease. The resulting protected fragment of RNA, now an sfRNA, functions as a genome-independent noncoding RNA (**Fig. 1A**). The ability of xrRNAs to block progression of the exoribonuclease is conferred by a complex and compact folded structure centered on a three-way junction and stabilized by multiple tertiary contacts (**Fig. 1B**) (41–44). The structure forms a ring-like motif through which the 5′ end of the resistant RNA element passes. Biochemical and biophysical analyses suggest that this novel topology acts as a molecular brace against the surface of the exonuclease which prevents it from progressing past a defined point (45–51). Structural and bioinformatic analysis strongly suggest that versions of this fold, and the resultant exoribonuclease-resistant function, are shared among all sfRNA-producing flaviviruses (44, 52). In addition, other classes of xrRNAs have been found in distal viral clades and thus far, the ring-like topology is a unique defining feature (44, 53–58).

**Figure 1.**
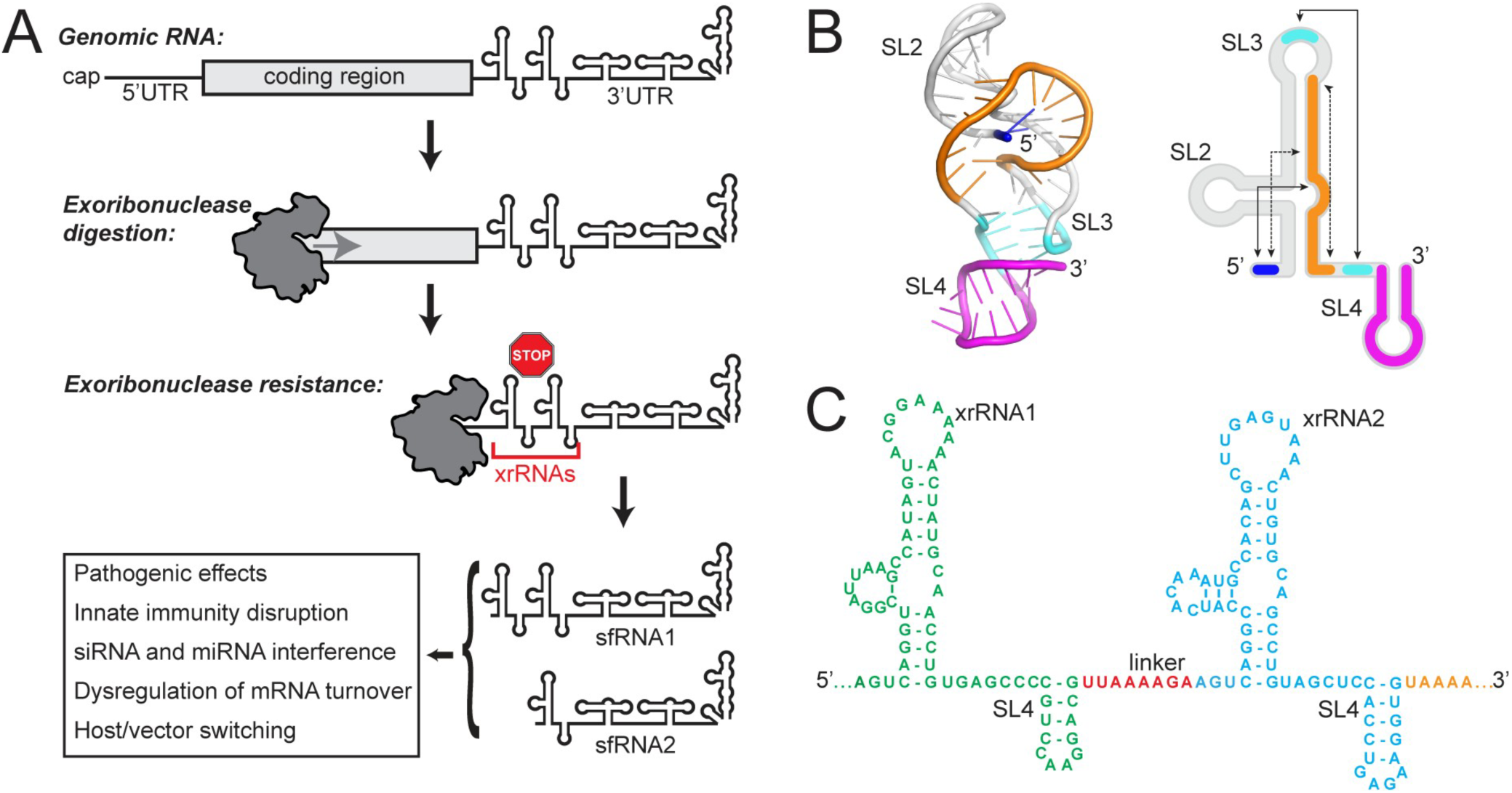
Orthoflavivirus sfRNA production by xrRNAs. (A) Cartoon representation of Xrn1 degradation of the flaviviral genome during infection, leading to sfRNA formation. (B) Three-dimensional structure of the first xrRNA from Zika virus (left) and secondary structure cartoon (right). The ring-like structure is shown in orange, the 5′ end in blue. An important pseudoknot interaction is shown in cyan, and a stem-loop (SL4) in magenta. Important long-range base pairs are indicated with arrows, a dashed line indicates a necessary base triple. (C) Secondary structure representation of the DENV2 tandem xrRNAs. xrRNA1 is green, xrRNA2 is cyan, the A-rich linker is red, RNA downstream of xrRNA2 is orange.

A striking feature of many flaviviruses, including DENV, is the presence of two or more consecutive xrRNA elements located within their 3′ UTR (**Fig. 1A&C**). When present, we refer to these as xrRNA1 and xrRNA2. These tandem xrRNAs are not identical in sequence and there are subtle differences in their characteristics (36, 59), although in all biochemically examined cases they can block Xrn1, and conserved sequences strongly indicate all can form a ring-like 3D topology (54). A simple hypothesis for the presence of tandem xrRNAs is that the functionally independent elements provide redundancy - any exoribonuclease that makes it through the first xrRNA is blocked by the second, ensuring sufficient sfRNAs are made. Indeed, in many viruses that have tandem xrRNAs, infection leads to two versions of sfRNA (**Fig. 1A**) resulting from the two xrRNAs (7, 33, 36).

The presence of tandem xrRNAs in many flaviviruses raises the question of why this arrangement is maintained. While the simplest explanation is redundancy, multiple lines of evidence suggest a more complex regulatory role. First, studies of several flaviviruses indicate that the function of tandem xrRNAs, when present, are coupled. That is, the structural integrity of one xrRNA affects the function of the other, suggesting some form of communication between the two seemingly independent structures (31, 32, 34, 42, 60). In DENV, WNV, and ZIKV infections, mutation of xrRNA2 to alter its structure and function reduces the accumulation of sfRNA2, as expected. However, these mutations also surprisingly markedly reduce the ability of xrRNA1 to block Xrn1, resulting in reduced sfRNA1 accumulation. Stated another way, this indicates that within tandem xrRNAs, the upstream xrRNA1 senses the presence or integrity of the downstream xrRNA2, altering xrRNA1’s function. This communication or coupling is observed in multiple flaviviruses, suggesting it is an evolutionarily conserved and maintained feature important to the fitness of many flaviviruses (13, 32, 36, 42, 60).

The unexpected coupling between tandem xrRNAs appears to be mechanistically linked to the specific generation of the sfRNA patterns required for some viruses to adapt to different hosts. Studies of DENV2 revealed that viral fitness in a mosquito vector vs. human host required different patterns of sfRNAs (31). Briefly, during infection in mosquitos, DENV2 accumulates mutations in xrRNA2 in sequences involved in critical tertiary structure interactions. These mutations expectedly eliminate sfRNA2 production while the coupling also reduces sfRNA1 produced by exoribonuclease halting at xrRNA1. This sfRNA pattern benefits the virus in mosquitos preferentially, but when virus harboring these mutations is used to infect human cells, the xrRNA2 mutations revert, resulting in the sfRNA pattern for efficient infection in humans and suppressing the immune response. Thus, in DENV2 xrRNA2 appears to be a key regulatory switch in which dynamic sequence changes affect the distinct but coupled xrRNAs (31), leading to changes in sfRNAs that allow the virus to switch between vector and hosts, maintaining and adjusting fitness as needed.

The coupling of tandem xrRNAs and resultant sfRNA patterns are important for DENV2 host-vector switching and fitness, but the effects appear to vary across different flaviviruses. For example, infection by ZIKV exhibits a different dependence on specific sfRNA species in host vs. vector, but the coupling effect between xrRNA1 and xrRNA2 remains evident (34). Interestingly, altering ZIKV xrRNA1 to eliminate sfRNA1 production, while still allowing sfRNA2 production, did not attenuate genomic replication in the liver but reduced genomic replication in the brain and placenta of mice (35). This cell type-dependent variation suggests that the specific pattern of sfRNAs, and thus perhaps coupling of the tandem xrRNAs, can also affect cellular tropism within a host. Insect-specific flaviviruses (ISFVs) have been identified with up to four xrRNAs in their 3′ untranslated regions but as these viruses do not switch hosts, the purpose of these multiple xrRNAs must differ from that in MBFVs. Indeed, a recent report suggests that multiple xrRNAs in ISFVs largely provide redundancy (61). Together, these data suggest variation in the role of tandem or multiple xrRNAs in different viral clades or species.

How tandem xrRNA coupling is achieved has been unanswered. Possibilities include a direct physical interaction between tandem xrRNAs within the same copy of viral genome, or bridging proteins, possibilities which are not mutually exclusive. Although high-resolution structures of individual xrRNAs have been solved (43, 54, 60), there is no similar structural information on tandem xrRNAs. Small-angle x-ray scattering experiments on tandem xrRNAs from several viruses yielded largely extended envelopes with no obvious interaction between the xrRNAs. However, as these are populationally weighted averages, they do not reveal individual conformations within a potentially dynamic structural ensemble that could provide insight into coupling (62).

To explore the sequence and structural requirements for xrRNA coupling, we used a combination of virology, molecular biology, biochemistry and structural biology. We focused on DENV2 as it represents a global health threat and the importance of coupling to host-vector switching is clear (31). Our data reveal that coupling can occur independent of viral infection and appears to be a characteristic of the RNA itself, operating identically in mammalian and mosquito cells. We find that the coupling effect depends on a conserved but unpaired A-rich RNA linker between the two xrRNA elements that may interact with an adjacent stem-loop to structurally bridge the two xrRNAs. Mutation of this linker leads to local structural destabilization and increased flexibility that then appears to alter the relative orientation and conformational dynamics of the tandem xrRNAs. Our results suggest new hypotheses for how this could lead to coupling of function during viral infection.

## RESULTS

### Mutations to DENV2 tandem xrRNAs alters sfRNA during infection

To explore the coupling between DENV2 tandem xrRNAs, we examined the pattern of sfRNA formation using wild-type (WT) DENV2 and various mutants to infect Vero cells (**Fig. 2A&B)**. With WT DENV2, Northern analysis revealed primarily accumulation of sfRNA1 and much smaller amounts of sfRNA2, formed from xrRNA1 and xrRNA2, respectively (**Fig. 2B**). We then tested mutant viruses in which the structural integrity of the xrRNAs was disrupted using a previously characterized C→G point mutation within the three-way junction (**Fig. 2B**) (42). Consistent with previous observations, destabilization of xrRNA1 led to a loss of sfRNA1 but essentially the same amount of sfRNA2 relative to WT. Disruption of xrRNA2 led to a loss of sfRNA2, but also a dramatic decrease in sfRNA1, illustrating the coupling effect. Viruses with mutations to both xrRNAs did not produce sfRNA1 or 2, as expected.

**Figure 2.**
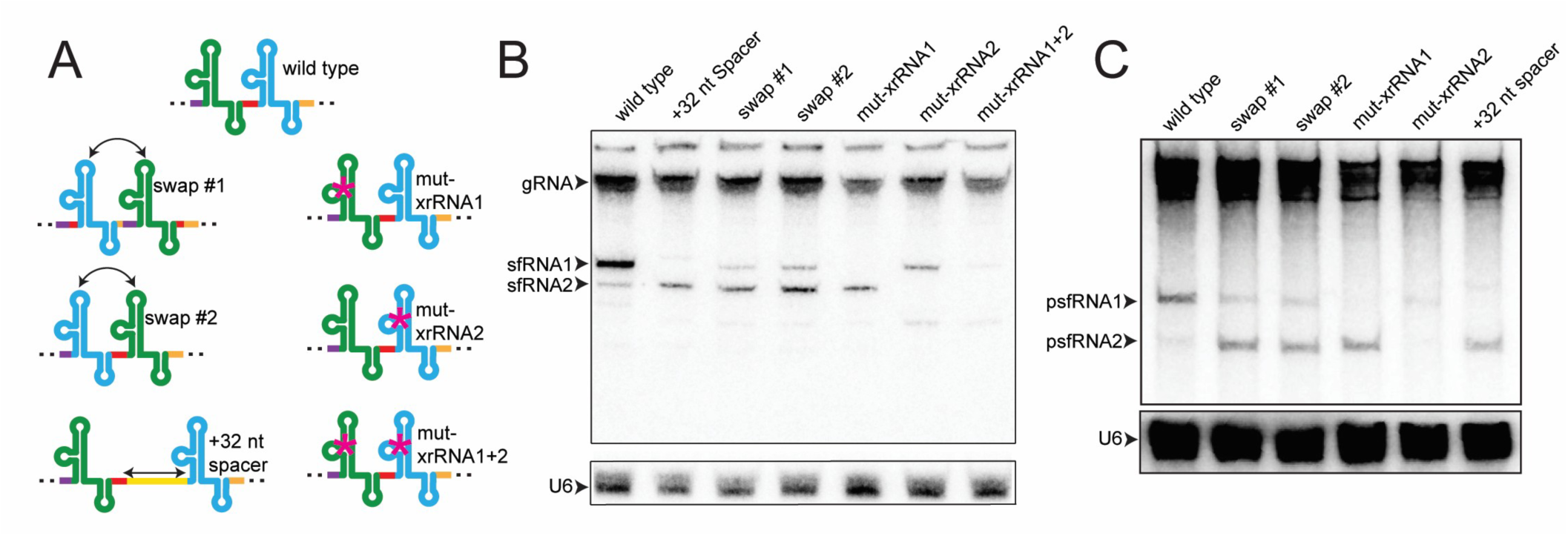
Effect of altering the tandem xrRNAs on RNA coupling. (A) Cartoon representations of the mutants used to test the coupling effect, colored as in Figure 1C. Full sequences and details are contained in Supplemental Figure S1. (B) Northern analysis of the sfRNA generated during infection of Vero cells with DENV2 wild type or viruses containing the changes in panel A, MOI = 1. Northern probe was complementary to a region downstream of xrRNA2. gRNA = genomic RNA. U6 was used as a loading control. (C) Northern analysis of the psfRNA generated in Vero cells with the NMD-based reporter system.

We next assessed if the coupling effect depends on the order of the two similar but not identical tandem DENV2 xrRNAs. Two versions of mutant DENV2 were created in which the positions of the xrRNAs were switched; the two versions differed by how much adjacent sequence was swapped along with the xrRNA structure (**Fig. 2A; Fig. S1**). Infection with either mutant lead to an altered sfRNA pattern relative to WT in which the amount of sfRNA1 was less than the amount of sfRNA2 (**Fig. 2B**). This result indicates that the order of the two xrRNAs matters, that they are not equivalent in their ability to halt Xrn1 during infection, and that xrRNA1’s ability to block Xrn1 decreases when it is not followed in the sequence by xrRNA2. These results are fully consistent with previous observations (31).

A possible explanation for the coupling effect is that the tandem xrRNAs directly physically interact. We tested this idea by introducing extra sequence between them, creating a long, flexible linker that allows for greater spatial distance between the xrRNA structures (“+32 nt spacer” **Fig. 2A**). This insertion mutant resulted in a pattern of sfRNA formation like that from mut-xrRNA1, indicating that if xrRNA2 is not directly downstream from xrRNA1, the latter’s ability to halt Xrn1 is altered (**Fig. 2B**). Put together, these results suggest that in DENV, xrRNA2 enhances xrRNA1’s ability to block Xrn1 in a mechanism that requires spatial proximity and is specific to the order of the xrRNAs.

### xrRNA coupling does not require viral infection

One possible mechanism for xrRNA coupling is the presence of a bridging viral protein or some other infection-specific condition. To create an experimental system to test this, we adapted a reporter system that was previously used to explore nonsense-mediated mRNA decay (NMD) (63). Briefly, this system uses a mammalian expression vector to express a β-globin mRNA containing a premature termination codon (PTC), followed by the 3′ UTR from DENV2 (**Fig. S2**). The PTC activates the NMD pathway, recruiting Xrn1 to the reporter RNA, which is then degraded until the xrRNAs are encountered, creating ‘pseudo-sfRNAs’ (psfRNAs) that are then visualized by Northern analysis.

We transfected Vero cells with the reporter containing the WT 3′ UTR from DENV2, resulting in a psfRNA pattern that closely resembles the sfRNA pattern from infection: predominantly psfRNA1 and much less psfRNA2 (**Fig. 2C**). Likewise, the psfRNA patterns resulting from reporters carrying mutations (mut-xrRNA1, mut-xrRNA2, swap #1, swap #2, +32 nt spacer) closely matched those observed during infection. Hence, the specific patterns of sfRNA produced by DENV2 tandem xrRNAs and the coupling between them does not require infection-specific proteins or conditions but does depend on specific characteristics of the RNA to include the structural integrity, proximity, and order of the two xrRNAs. In addition, because this NMD-based reporter system faithfully recapitulated observations obtained from infection models, it provides a simple way to test additional mutants without generating mutant virus or infecting cells.

### Sequence conservation reveals an A-rich linker between tandem xrRNAs

A possible mechanism for the coupling between tandem xrRNAs in DENV2 is a direct physical interaction between the two xrRNAs. As coupling has been observed in tandem xrRNAs from multiple species, this interaction could be facilitated by conserved sequence and structural features. These features might not be conserved within individual xrRNAs but unique to those positioned in a tandem arrangement. To explore this, we aligned the region comprising xrRNA1, the intervening linker, and xrRNA2 from multiple MBFVs based on primary sequence and secondary structure guided by previous 3D structural data and alignments (43, 54, 60) (**Fig. 3A**).

**Figure 3.**
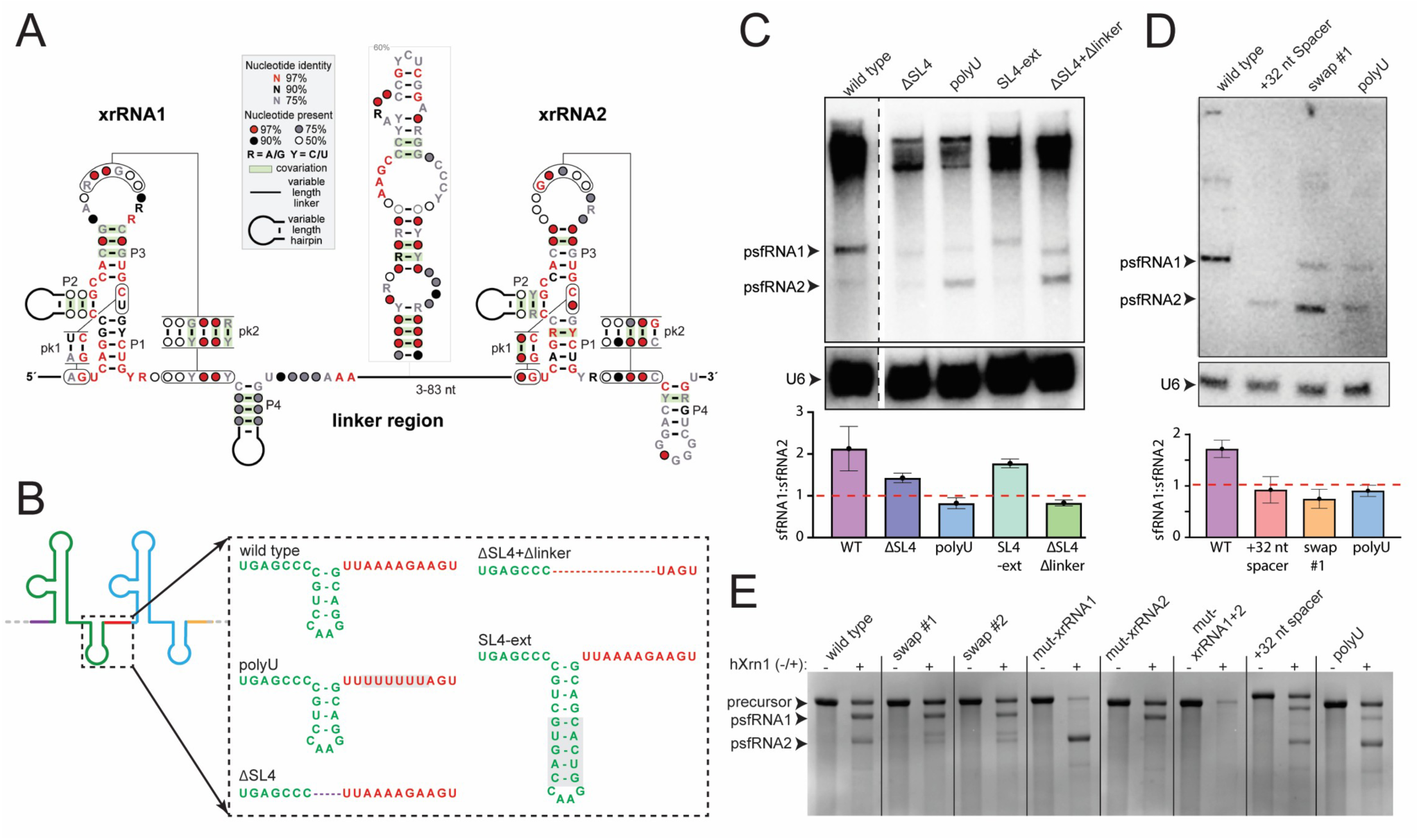
Mutations to the SL4 and the A-rich linker region disrupt psfRNA patterns. (A) Consensus secondary structure model of tandem xrRNAs created by R-Scape, using the alignment shown in Figure S2. (B) 2D cartoon representation of the tandem xrRNAs showing the mutants tested in panel C and D. Shading in the SL4-ext mutant indicates inserted base pairs. (C) Northern analysis of the psfRNA generated using the mammalian NMD-based reporter system (**Fig. S1**). A graph of the psfRNA1:psfRNA2 ratio is shown below the blot, the red dashed line indicates a ratio of 1. The probe targeted sequence downstream of xrRNA2. U6 served as a control. (D) Northern analysis of the psfRNA generated with the insect NMD-based system (**Fig. S1)**. A graph of the psfRNA1:psfRNA2 ratio is shown below the blot, the red dashed line indicates a ratio of 1. The probe targeted sequence downstream of xrRNA2. U6 served as a control. (E) Stained gel of the analysis of *in vitro* degradation of mutant tandem xrRNAs by purified hXrn1. Precursor and psfRNAs generated by partial digestion are indicated.

Examination of the consensus sequence and secondary structure model did not reveal any nucleotide positions likely to participate in any obvious base pairing or tertiary interactions between the two xrRNAs (**Fig. 3A**). Indeed, most areas of conservation are consistent with sequence-dependent secondary and tertiary interactions within each xrRNA known to be essential for exoribonuclease halting, with one exception. We noted a set of conserved adenosine nucleotides in the region that links the two xrRNAs (**Fig. 3A**). Further examination of individual viral sequences revealed that most contain at least three consecutive adenosine nucleotides, and many contain additional adenosines upstream or downstream of this (**Fig. S3**). This is true both when an intervening stem-loop is present between the two xrRNAs and when it is absent. In DENV2, this ‘A-rich linker’ sequence is 5′-UUAAAAGA-3′. Importantly, this region and the stem-loop that precedes it (SL4) are not necessary for halting exoribonuclease activity and there is no evidence that the A-rich region is involved in any base pairing. Nonetheless, examination of previously published chemical probing data show that it is well protected from modification, suggesting it adopts a stable structure in the context of tandem xrRNAs (42). Adenosines readily stack and often form RNA tertiary interactions (64–67), so we hypothesized that this A-rich linker region may be important for the coupling between tandem xrRNAs in DENV2.

### The A-rich linker region is necessary for tandem xrRNA coupling

To test the hypothesis that the A-rich linker is important for xrRNA coupling, we made a series of mutants targeting this region and tested them using the NMD-based reporter system (**Fig. 3B**). Because the absolute amount of psfRNAs produced was somewhat variable (likely due to variation in transfection efficiency), changes in sfRNA patterns were measured and reported as the ratio of psfRNA1 to psfRNA2.

With WT DENV2 in mammalian cells, approximately 2X more psfRNA1 is produced than psfRNA2, which changes when mutations are introduced (**Fig. 3C**). Removing SL4 but maintaining the A-rich sequence (ΔSL4) or increasing the size by adding more base pairs to its stem (SL4-ext) both resulted in a decrease in the amount of psfRNA 1 relative to psfRNA 2. However, in both cases the psfRNA1:psfRNA2 ratio remained above 1. More dramatic changes in psfRNA patterns were observed when the A-rich linker was altered. Deletion of SL4 and the A-rich region together (ΔSL4 + Δlinker) resulted in a switch in which less psfRNA1 was produced relative to psfRNA2. Focusing on the A-rich linker, substituting the A-rich region with a stretch of U bases (polyU) likewise resulted in a psfRNA1:psfRNA2 ratio below 1. Of note, this switch in relative abundances of the psfRNAs is similar to that observed when the xrRNA2 is destabilized or the two sfRNAs are spatially separated (**Fig. 2C**). Importantly, in the polyU mutant neither xrRNA has been mutated and no predicted base pairs are broken. We conclude that the A-rich linker is necessary for DENV2 tandem xrRNA coupling in mammalian cells and likely works together with SL4.

### The A-rich linker is important for coupling in mosquito cells

Given the differences in sfRNA patterns in mosquito versus mammalian cell infection models, we asked if the A-rich linker is critical for coupling during mosquito infection. We first generated a surrogate reporter system to work in C6/36 cells by again adapting the NMD-based reporter (**Fig. S2**). Specifically, we used an endogenous mosquito alcohol dehydrogenase gene containing a PTC in the second exon to trigger Xrn1 degradation of the xrRNAs (68). This system was again able to recapitulate the wild type and mutation-induced changes in psfRNA patterns observed during infection, indicating that it can reliably report on how mutations alter sfRNA patterns in mosquito cells (**Fig. 3D**).

Using this reporter in mosquito cells yielded a pattern with more psfRNA1 relative to psfRNA2, similar to what we observe in mammalian cells. Likewise, the +32 nt spacer and swap #1 mutants resulted in a switch of the relative psfRNA1 to psfRNA2 abundance. Altering the A-rich linker to the polyU sequence resulted slightly more psfRNA2 than psfRNA1 (**Fig. 3D**). Thus, these mutants result in a switch in the relative abundances of the two psfRNAs in both mammalian and mosquito cells, indicating that the coupling effect is occurring and conserved in both host and vector cells. These results agree with previous reports that sfRNA patterns and host adaptation do not depend on differences in the host machinery (31).

### The A-rich linker is important for coupling in reconstituted resistance assays

As the functional coupling between tandem DENV2 xrRNAs does not depend on viral infection or species-specific cellular machinery, we tested the hypothesis that coupling is conferred by characteristics of the RNA itself and its interactions with Xrn1. We used an established *in vitro* assay in which both WT and mutant tandem DENV2 xrRNAs were challenged with purified human Xrn1 (hXrn1) to assess their ability to block progression of the enzyme, the presence of coupling, and the effect of mutations on both. Under our experimental conditions, treatment of WT RNA by hXrn1 resulted in two degradation-resistant RNAs, consistent with the function of the two tandem xrRNAs (**Fig. 3E**). Assignment of these resistant products was confirmed by point mutations to either xrRNA1 or xrRNA2 (mut-xrRNA1, mut-xrRNA2), which resulted in a loss of the corresponding gel band, and mutation of both led to the expected loss of both products (mut-xrRNA1+2).

To determine if mutants that affect coupling in cells also affect it *in vitro*, we then tested the mutants in which the two xrRNAs were swapped, the +32 spacer mutant, and the polyU mutant (**Fig. 3E**). For both swap mutants, we observed a change in the amount and pattern of the smaller resistant product, but not the same effects as were observed in cells with these mutants. Likewise, the +32 nt spacer appeared to result in a slight increase in the smaller product relative to the larger, but a less obvious effect than was observed in cells. In contrast, hXrn1 treatment of the polyU mutant resulted in a clear change in the pattern of resistant products, with less of the larger xrRNA1-dependent product and relatively more of the xrRNA2-dependent product (**Fig. 3E**). This pattern is similar to what was observed with the mutant in reporter assays in both mammalian and mosquito cells (**Fig. 3C&D**).

Together, these results indicate that the biochemical conditions of the *in vitro* assays do not fully recapitulate the loss of coupling by all mutants – perhaps the effects of these mutants are too subtle and the *in vitro* assay is not sufficiently stringent, or an unidentified cellular protein plays a role. Nonetheless, the polyU mutant exhibits a clear loss of coupling in the *in vitro* assay, consistent with the conclusion that the sequence of the unpaired A-rich linker region between the two DENV2 tandem xrRNAs is necessary for coupling and its effect is inherent to the RNA.

### Mutation of the A-rich linker results in local structural changes

Insertion of sequence between the two xrRNAs and mutation of the A-rich linker each alter DENV2 xrRNA coupling, so we therefore asked what the effect of these mutations are on structure. Do the mutations cause a loss of long-range interactions or a large change in secondary structure? To assess how these mutants alter RNA secondary and perhaps tertiary interactions, we utilized selective 2′-hydroxyl acylation analyzed by primer extension and mutational profiling (SHAPE-MaP) chemical probing (69, 70).

The chemical probing profile of the WT RNA shows a pattern of modification consistent with the known xrRNA secondary structure and tertiary interactions (**Fig. 4A**). Germane to this study, the A-rich linker, despite being predicted to not be involved in any base pairing, shows more protection from modification than would be expected from a series of unpaired bases. This indicates a lack of local conformational flexibility, suggesting the linker is involved in some sort of stable structure. In contrast, in the +32 nt spacer mutant the extended linker is highly modified, indicating it is conformationally flexible (**Fig. 4B**). However, in the +32 nt linker the chemical probing of each xrRNA remains virtually identical to WT, and the A-rich linker that follows xrRNA1 also maintains a similar chemical probing pattern as in WT. We interpret these observations as indicating that the two xrRNAs are properly folded and the A-rich linker is still involved in a local stable structure despite the insertion of a linker between it and xrRNA2.

**Figure 4.**
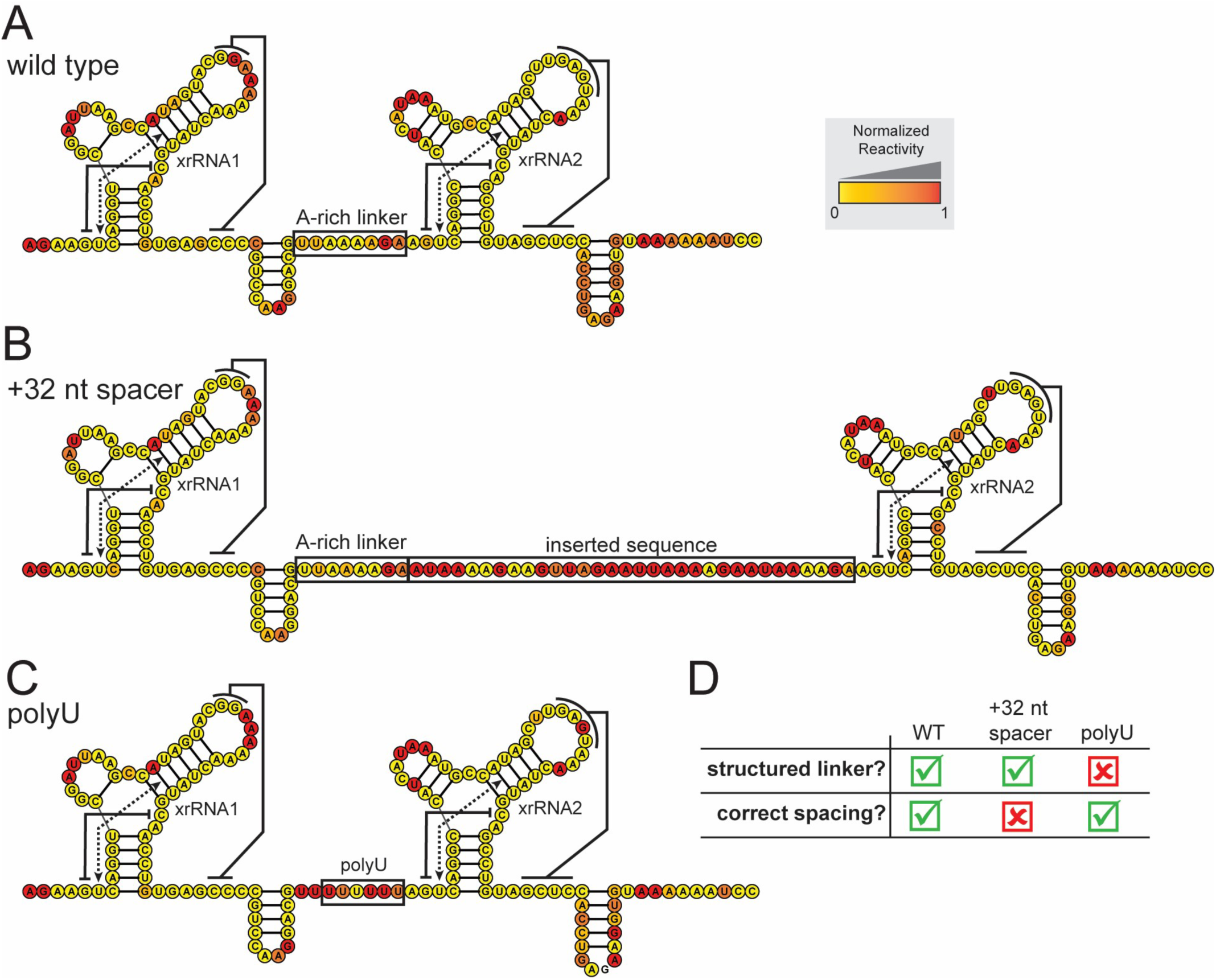
SHAPE-MaP analysis of WT and mutant DENV2 tandem xrRNAs. (A) Results of SHAPE chemical probing mapped onto the secondary structure diagram of the WT DENV2 tandem xrRNAs. Key tertiary interactions and structural elements are labeled. (B) Chemical probing results mapped onto the secondary structure diagram of the +32 nt spacer mutant DENV2 tandem xrRNAs. (C) Chemical probing results mapped onto the secondary structure diagram of the polyU mutant DENV2 tandem xrRNAs. (D) Table illustrating how the two mutants each lack one of the key requirements for tandem xrRNA coupling, reflected in the probing.

We then probed the polyU mutant (**Fig. 4C**). As with the WT and the +32 nt spacer RNA, there was no significant change in the modification of the two xrRNAs relative to WT, indicating that each was individually folding correctly. However, the polyU mutation resulted in a clear increase in modification in the linker region, indicating a marked change in local conformational flexibility. Importantly, this increase was limited to the linker, indicating that the changes in flexibility are localized to this region of structure and are unlikely to involve changes in base pairing.

Contextualizing the chemical probing results with the Northern analysis leads to several conclusions. First, coupling and the pattern of DENV2 sfRNA formation require correct folding of each individual xrRNA, but this is not sufficient. Second, coupling depends on an A-rich linker-dependent local structure between the two xrRNAs, which is lost in the polyU mutant (**Fig. 4D**). Third, coupling and proper sfRNA biogenesis requires that the two folded xrRNAs be in spatial proximity, which is broken in the +32 nt spacer mutant (**Fig. 4D**).

Interestingly, when we probed an RNA in which the positions of the two xrRNAs are switched (swap #1), we see little or no change in the chemical probing pattern in any part of the RNA (**Fig S3**). Thus, it appears that in this mutant, the A-rich region is structured and can still form putative tertiary contacts as WT, but the resultant structure is not able to productively couple the two xrRNAs. This suggests that the coupling effect depends on the A-rich linker, but that there are subtle differences between nonidentical xrRNA1 and xrRNA2 that are also important.

### CryoEM analysis of tandem xrRNAs reveals a bridging linker-containing structure

The chemical probing and functional data suggest that local structure involving the A-rich linker is essential for DENV2 tandem xrRNA coupling, raising the question of how this feature connects the two xrRNAs. Although the structures of individual MBFVs xrRNAs have been solved by crystallography, no such structure exists for tandem xrRNAs. We therefore used single-particle cryo-electron microscopy (cryoEM) to interrogate the structure of the WT DENV2 tandem xrRNAs (**Fig. 5, Fig. S5**). Analysis of the resultant particles revealed a heterogenous sample, but one conformational state emerged that allowed for the reconstruction of mid-resolution maps of the RNA (**Fig. S4**). No other well-populated state was found. While maps at this resolution could not inform on fine structural details or allow *ab initio* building of nucleotide-resolution structures, they were sufficient to allow modeling of the DENV2 tandem xrRNA architecture and mode of interaction.

**Figure 5.**
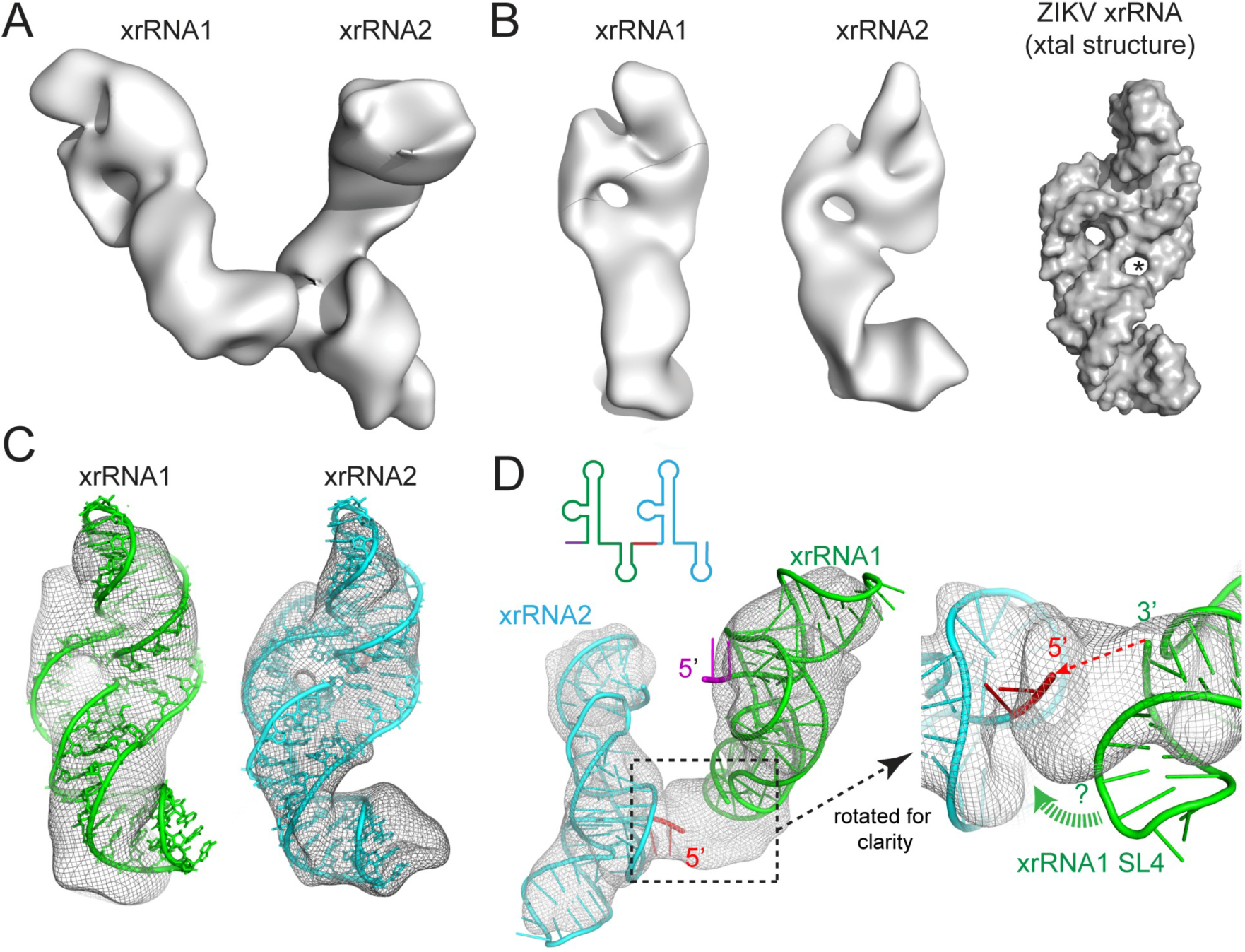
CryoEM analysis of the DENV2 tandem xrRNAs. (A) Single-particle analysis cryoEM map of the tandem xrRNAs from DENV2 shown as a surface representation. Details of the analysis are found in Figure S4. (B) The two lobes from the map shown separately. The lobe that contains xrRNA1 is on the left, xrRNA2 in the middle. A surface representation of an xrRNA from ZIKV solved by x-ray crystallography is shown to the right for comparison. (C) Docking of the ZIKV xrRNA structure into the two lobes of the DENV2 tandem xrRNA cryoEM map. The structure at the top extends out of the map by a base pair, consistent with the different sizes of the SL2 of ZIKV and DENV2. (D). Two docked xrRNAs in the map, colored to match the secondary structure cartoon. Right: closeup of the region between the two xrRNAs. The 3′ end of xrRNA1 is placed to connect to the 5′ end of xrRNA2 by the A-rich linker. SL4 would need to swing into a new position (green dashed arrow) to properly fit the map. This would place the SL4 minor groove adjacent to the A-rich linker.

Globally, the cryoEM maps comprise a two-lobed structure. The two lobes are similar in shape, are held at a specific relative orientation, and are linked through a single bridging feature (**Fig. 5A**). Consistent with the two linked elements being the two tandem xrRNAs, examination reveals that each contain a ring-like feature matching other xrRNA structures (**Fig. 5B**). To model these two map elements as xrRNAs, we docked the xrRNA1 crystal structure from ZIKV into each of the two cryoEM map elements. ZIKV serves as a reasonable model as it is similar in sequence and secondary structure to the DENV2 xrRNAs. Docking of this ZIKV xrRNA revealed remarkably good agreement between the structure and the map, to include the structure and location of the ring-like feature and in some places the presence of major and minor helical grooves (**Fig. 5C**). Several of the differences between the map and docked xrRNAs matched differences in the length of helical elements between DENV2 and ZIKV xrRNAs, further validating the map and the global placement of the modeled tandem xrRNAs. Docking of the xrRNA structures into the map immediately indicated which was xrRNA1 and which was xrRNA2, as the 3′ end of the former aligned with the 5′ end of the latter (**Fig. 5D**). Notably, the overall fit of the structure into the xrRNA2 map feature was noticeably better than into the xrRNA1 map feature (**Fig. 5C**).

The cryoEM map and resultant model show that in this configuration, the two xrRNAs are not packed against one another, but instead adopt a specific orientation connected through a single map feature (**Fig. 4D**). This feature corresponds to the sequence between the two xRNAs; that is, SL4 and the A-rich linker downstream of xrRNA1. Notably, however, this region is the only part where the map and docked xrRNA1 deviate substantially from one another. Specifically, in the crystal structure of the ZIKV xrRNA1, SL4 stacks roughly coaxially on the base pairs that comprise pseudoknot 2 (**Fig. 1B**), while the map feature we assign to SL4 is positioned at an angle pointed towards the 5′ side of xrRNA2. The map is strong in this region and appears large enough to contain both SL4 and the A-rich linker, although the resolution is not sufficient to distinguish these elements or place nucleotides. However, the cryoEM map and model strongly suggest that SL4 and the A-rich linker together form a specific structure that bridges the two xrRNAs.

### Local structural disruption in the A-rich linker alters global tandem xrRNA conformation

Our combined findings indicate that the A-rich linker is necessary for DENV2 tandem xrRNA coupling, and the cryoEM maps suggest that this linker and SL4 form a structural bridge between the two xrRNAs. A logical prediction is that mutations to the A-rich linker disrupt this bridge and the resultant alteration of local structure would lead to a change in global conformation. To test this, we used size exclusion chromatography (SEC) coupled with small-angle x-ray scattering (SAXS) to interrogate the global solution conformations of WT and mutant DENV2 tandem xrRNAs at several magnesium concentrations. The use of in-line SEC helped ensure a monomeric and monodisperse sample.

SEC-SAXS data were collected on WT DENV2 tandem xrRNAs at 0, 2, and 4 mM MgCl2 and analyzed to yield information about the solution biophysical properties of the RNA. Examination of the Kratky plots for these data indicate folding of the RNA upon the addition of magnesium (**Fig. 6A**). Folding appears complete at 2 mM Mg^2+^, as additional Mg^2+^ to 4 mM did not result in additional changes. Likewise, the pairwise distance distribution function graph for WT RNA shows folding at 2 mM Mg^2+^, with a decrease in the Dmax accompanying the folding event and no additional decrease with more Mg^2+^. The overall shape of the curve suggests a conformation comprising two separate lobes linked by a thinner connecting structure. This is consistent with the cryoEM map of the tandem xrRNA architecture, with the two xrRNAs comprising two larger domains linked by a thinner bridging feature (**Fig. 5A**). Taken together, these data indicate a magnesium ion-induced stabilization of the 3D fold of the RNA, demonstrating that SEC-SAXS is suitable to monitor the biophysical characteristics of the global tandem xrRNA conformation.

**Figure 6.**
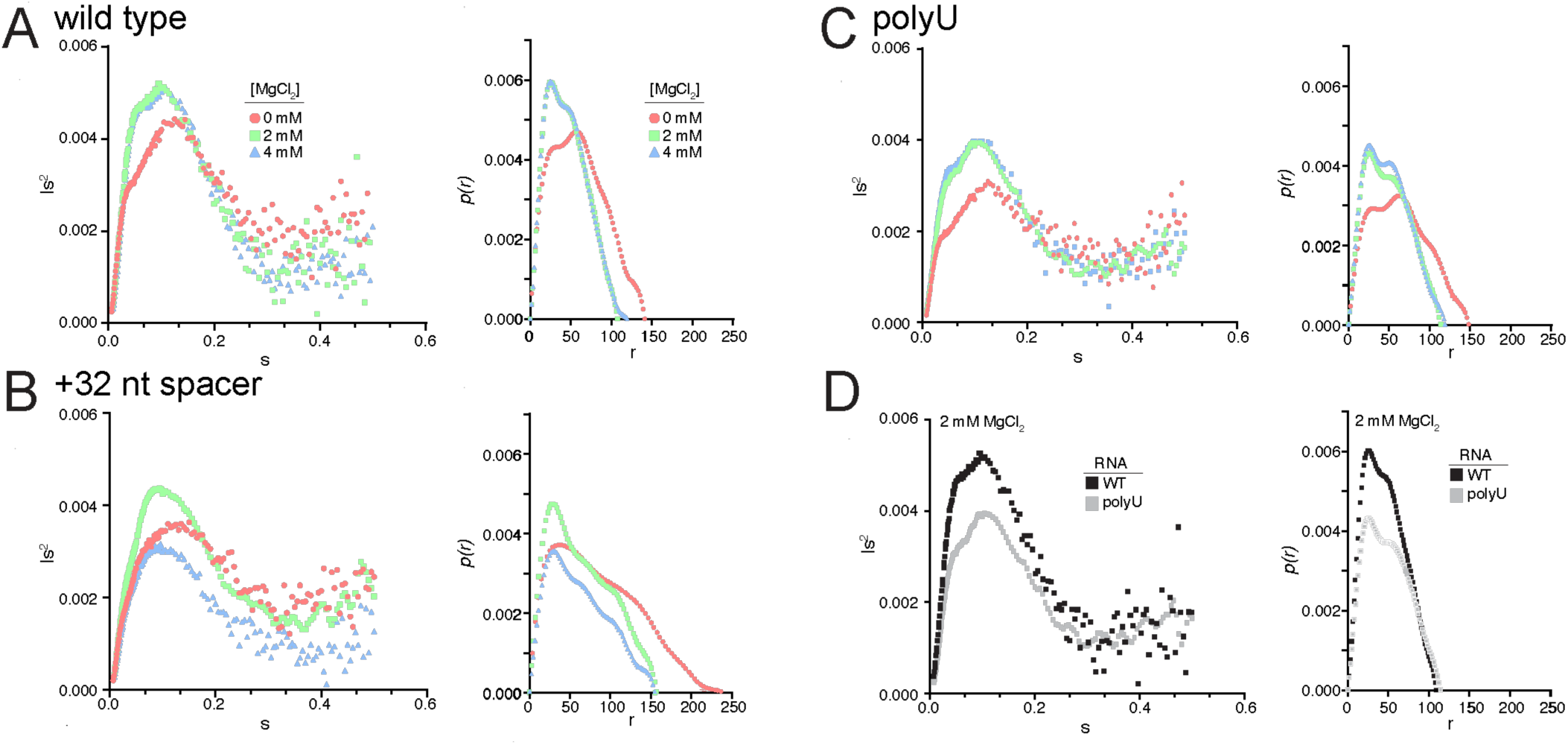
SAXS analysis WT and mutant DENV2 tandem xrRNAs. (A-C) Kratky plot (left) and distance distribution function plot (right) for each of the tandem xrRNA constructs, comparing data obtained in the presence of 0, 2, or 4 mM Mg^2+^. (D) Kratky plot (left) and distance distribution function plot (right) comparing WT and polyU at 2 mM Mg^2+^ concentration. Results from the swap #1 mutant are in Figure S6.

We next examined the +32 nt spacer mutant by SEC-SAXS, both to verify that spatially separating the two xrRNAs with intervening sequence would result in a change in the measured solution biophysical properties, and to provide a rough calibration of the degree of the observed change. As with WT, the Kratky plots for the +32 nt spacer showed a response to the addition of 2 mM Mg^2+^ but the curve differed dramatically from WT, indicating a more extended structure (**Fig. 6B**). This is also observed in the pairwise distance distribution function again and a larger Dmax value. This is consistent with the 32 nt sequence insertion producing an overall larger and more extended conformation with the two xrRNAs being more widely separated (**Fig. 6B**). In addition, while the WT RNA showed no additional changes with more Mg^2+^, addition of more Mg^2+^ to the +32 nt spacer RNA resulted in additional structural changes, suggesting less stable interactions can be promoted at higher cation concentrations. Overall, these data combined with the chemical probing data and functional data are consistent with the +32 nt spacer RNA forming a more extended structure in which the two DENV2 xrRNAs are no longer in proximity due to their connection by a flexible and more extended linker. This change in global structure of the DENV2 tandem xrRNA conformation correlates with the disrupted tandem xrRNA coupling observed in the in cell and *in vitro* assays.

Having verified the ability of SEC-SAXS to report on global conformational differences in WT and mutant tandem xrRNAs, we subjected the polyU mutant to this analysis. The Kratky plot once again indicated a folding event with the addition of 2 mM Mg^2+^, but the plot again shows clear differences compared to WT (**Fig. 6C&D**). These differences are consistent with the two aforementioned lobes being less well ordered relative to each other or more distant from one another. Likewise, the pairwise distance distribution function plot shows a folding event induced by magnesium but a different shape and larger Dmax than WT (**Fig. 6C&D**). This is consistent with a slightly less compact or well-ordered structure. Lastly, as with the +32 nt spacer mutant, the addition of more magnesium to 4 mM resulted in additional changes to the structure, although this was less dramatic than for the +32 nt spacer (**Figs. 6C**). These data show that mutation of the unpaired A-rich linker is sufficient to alter the global conformational state in a way that is consistent with a change in the spatial relationship of the two tandem xrRNAs. Combined with the other data presented here, this is consistent with a model in which the A-rich linker is part of a structural bridge that links the two tandem xrRNAs, creating a specific orientation, leading to a coupling of function by a mechanism that remains to be fully understood.

## DISCUSSION

Many mosquito-borne flaviviruses contain tandem xrRNAs, and in DENV2 the exonuclease resistance function of the two is coupled. This coupling is important for controlling the type and abundance of sfRNAs generated during infection, influencing the ability of the virus to successfully infect different hosts (31–34, 36). However, the features of the tandem DENV2 xrRNAs that create this functional coupling have remained unknown. In this study, we used observations gleaned from infection models to motivate *in vitro* experiments, which allowed us to examine the inherent properties of the tandem DENV2 xrRNAs without confounding cellular variables. Specifically, we used a combination of virology, molecular biology, bioinformatics, biochemistry, and biophysics to identify the sequence and structural determinants of the coupling phenomenon. Our findings show a previously unrecognized A-rich linker sequence located between the two xrRNAs is necessary for coupling, likely interacting with an adjacent stem-loop structure to structurally bridge the two xrRNAs and position them relative to one another. These findings invite new hypotheses for how the resultant global architecture couples the DENV2 tandem xrRNAs.

Despite our discovery that the A-rich linker connecting the DENV2 tandem xrRNAs is essential for coupling, there is no evidence that this sequence forms any base pairs with the rest of the DENV2 3′ UTR. While seemingly counterintuitive, it is well established that poly(A) sequences form helical structures without pairing partners, as a single strand of adenosines readily stacks in an A-form geometry (71–73). Such structures are stable enough that they are only weakly modified in SHAPE chemical probing experiments that detect local backbone flexibility (74), and stretches of adenosines are readily recognized by proteins based on their 3D structure (75–77). Thus, the presence of an A-rich linker that is structured in the absence of base pairing partners, and that physically connects the two xrRNAs, is consistent with the known behavior of stretches of adenosines.

The A-rich linker is essential for coupling, and mutation of the sequence leads to the largest loss of coupling that we observed among our mutants. However, the adjacent SL4 may also play an important role, as the cryoEM maps and SAXS data suggest the linker and SL4 interact to create a structural bridge between the two xrRNAs. Although the map is not of sufficient resolution to confidently assign the position of individual nucleotides, they suggest how the interaction between these components could occur. If SL4 were to swing from its modeled position, it could readily occupy the map feature that links the two xrRNAs (**Fig. 5D**). In this proposed position, the minor groove of SL4 would align with the A-rich linker, allowing for the formation of tertiary contacts between the adenosines and the minor groove. A-minor interactions, in which the A base forms hydrogen bonds with functional groups in the minor groove, are common in RNA structure and have been observed in a wide variety of contexts (5, 64, 65, 78). In DENV2, A-minor interactions could stabilize the position of SL4 within a structure that spatially connects and bridges the two xrRNAs. The cryoEM map also suggests that the position of SL4 in the bridge places the apical loop of the SL to interact with the 5′ end of xrRNA2, but we cannot confidently predict the nature of these interactions.

Our findings strongly support the conclusion that the A-rich linker + SL4 create a bridge between the two xrRNAs, providing a physical connection. This now suggests a new question: how does this structural bridge functionally couple the two xrRNAs? That is, how is the structural integrity of xrRNA2 communicated to xrRNA1 through this bridge? Several not mutually exclusive hypotheses can be proposed.

First, the structural bridge between the two might thermodynamically couple the two xrRNAs. Specifically, the bridge transforms two loosely connected xrRNA structures, which are folded independently, into a single structural unit in which destabilization of one element results in an overall destabilization. One can imagine that disruption of xrRNA2’s fold would destabilize or alter the A-rich linker + SL4 bridge, which could then affect the behavior of xrRNA1. We note that although the cryoEM maps were of sufficient quality to allow unambiguous placement of each xrRNA, the fit of xrRNA2 was better than for xrRNA1. One intriguing possibility is that the repositioning of SL4 to interact with the A-rich linker and xrRNA2 could distort the structure of xrRNA1 in such a way as to enhance its ribonuclease resistance. This remains speculative, providing questions for ongoing exploration.

Another interesting possibility comes from the observation that the A-rich linker + SL4 structural bridge gives the two xrRNAs a preferred relative orientation (**Fig. 5**). It seems unlikely that this orientation is static but rather exists within a conformational ensemble. However, this conformation was the only one to emerge from the cryoEM analysis and the resultant map was easily interpreted with features consistent with the biochemistry and virology, suggesting it represents a thermodynamically preferred position. Consistent with this, mutation to the A-rich linker that disrupts only local structure (**Fig. 4**) results in a loss of coupling that the SAXS data shows changes the spatial relationship of the two xrRNAs (**Fig. 6**).

The relative orientation of the two xrRNAs is correlated to coupling through the SAXS analysis, but this does not strictly establish cause and effect or a mechanism by which this orientation would induce coupling. However, we note that in the observed orientation, an exoribonuclease approaching xrRNA1 from the 5′ direction would likely interact simultaneously with both xrRNA1 and xrRNA2, a possibility that is not apparent from the 2D drawings of tandem xrRNAs. Indeed, using the cryoEM map to build a model in which Xrn1 is placed on the tandem xrRNAs approximately where it would be when halted predicts interactions between Xrn1 and both xrRNAs (**Fig. 7**). In this speculative model, simultaneous interactions between xrRNA1, xrRNA2 and Xrn1 would enhance the ability of xrRNA1 to halt the enzyme, and mutation of the linker to disrupt the relative orientations of the xrRNAs would disfavor these interactions. This compelling idea suggested by our data remains untested.

**Figure 7.**
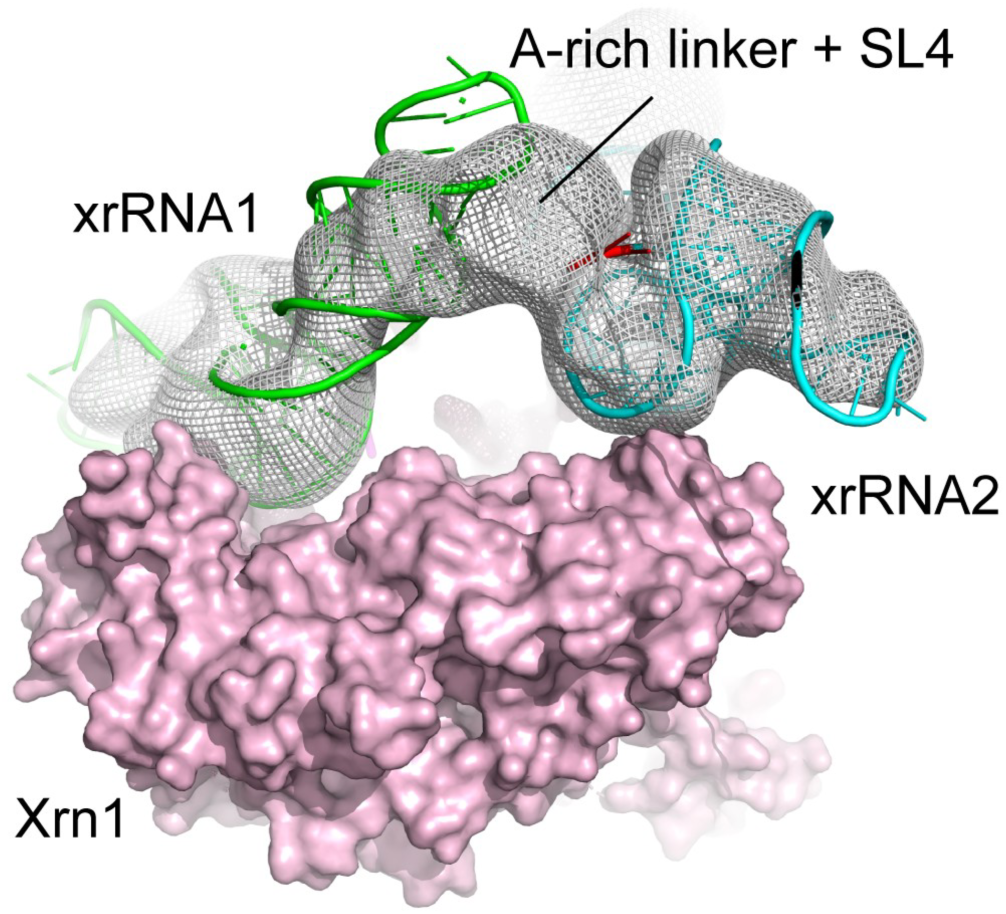
Model of Xrn1 on the tandem xrRNAs. The structure of Xrn1 (PDB entry 2Y35), solved by crystallography (86), was manually placed such that the 5′ end of xrRNA1 is positioned to enter the active site. The relative orientation of the two xrRNAs observed in the cryoEM maps places xrRNA2 such that it could interact with Xrn1 and xrRNA1 simultaneously.

In addition to disrupting the A-rich linker + SL4 bridge, we observed that swapping the two xrRNAs changed the relative abundance of the two sfRNAs, further supporting the notion that the two xrRNAs are not fully independent or identical elements but operate within an ordered structural unit where their relationship matters. One possible explanation for the changes in sfRNA patterns due to xrRNA position swapping is found in the fact that when the xrRNAs were swapped, the SL4s were included in the swap. We note that the length and apical loop of the SL4 from xrRNA2 is different from the SL4 from xrRNA1 – the former may not be able to substitute for the latter to form the stable bridging structure.

In this study, we limited ourselves to an examination of the tandem xrRNAs from DENV2, but tandem xrRNAs are common in mosquito-borne flaviviruses. However, the coupling effect does not exist identically in all of them or has not been examined in most species. The A-rich linker and adjacent SL4 are conserved elements but may not operate identically in all viral species or clades. Small changes to sequence, the size of SL4, or the 5′ side of xrRNA2 could alter tertiary contacts and key interactions and change the degree to which the xrRNAs are coupled, if at all. In addition, while our studies focused on the tandem xrRNAs from DENV2, which do not have an intervening SLIII sequence between them, many other viral species have this structural element between their tandem xrRNAs (**Fig. 3A, Fig. S3**), which could also play a role in coupling. Thinking broadly about viral evolution, these features may allow the coupling phenomenon to be adapted easily to the needs of species viral species in specific hosts, without changing the sequences, fold or function of the individual xrRNAs.

In summary, we have identified that a conserved A-rich linker is a necessary feature for coupling the tandem xrRNAs in DENV2, where it is part of forming a structural bridge between the two xrRNAs that likely includes xrRNA1’s SL4. Integrity of this bridge is important for regulating the relative abundance of the sfRNAs produced from these RNAs. These results provide an RNA structure-based explanation for observations for changing sfRNA abundance in different hosts during DENV2 infection and suggest new testable mechanistic hypotheses to guide further exploration.

## METHODS

### Dengue virus mutant generation

An AgeI restriction site was introduced into the DENV2 3′-subclone (pD2i-3′ABX) using the Q5 Site-Directed Mutagenesis Kit (NEB) according to the manufacturer’s instructions. Non-overlapping primers with the mutagenesis region on the forward primer only were designed with the recommended NEBaseChanger web interface (https://nebasechanger.neb.com). PCR mutagenesis and amplification of ∼23 ng of initial plasmid template following standard thermocycling conditions was confirmed via 0.5% agarose gel electrophoresis prior to performing the kit’s Kinase-Ligase-Dpn1 (KLD) reaction and transformation into DH5α competent *E. coli*. Briefly, 5 μL of KLD reaction was added to 50 μL competent cells on ice, incubated for 30 minutes prior to 40 second heat shock at 42°C. Transformation tube was returned to ice for 5 minutes before the addition of 950 μL LB broth and shaking at 250 rpm and 37°C for 45 minutes. 100 μL of bacteria were plated onto LB Agar - Carbenicillin (100 ug/mL) plates and incubated for 18-24 hours at 37°C. Small & clear colonies were chosen for mini-prep (5 mL LB, shaken at 225 rpm for 18 hours at 37°C) amplification of the plasmid, then cleanup according to manufacturer’s instructions (Qiagen 27104). Confirmation of correct insertion of AgeI site, as well as maintenance of the singular BspEI required for downstream molecular cloning processes, was conducted via sequential digest of 0.2-0.5 μg plasmid DNA with both restriction enzymes (AgeI-HF- NEB R3352L, BspEI- NEB R0540L), as well as Sanger sequencing of the mutagenized region with a primer complementary to the NS5 region of the genome (Sequence: 5′-CCAAGCAGCAATAAATCAAGTTAGATCCCTTATAG-3′). Finally, newly created infectious clone plasmid pD2i-3′△AgeIWT was re-transformed as above for maxi-prep (1 L) amplification of working stocks. To generate the virus 3′ UTR mutants, gBlocks (double-stranded DNA gene fragments) with the entire wild type (WT) or mutant 3′UTR of DENV2 were designed with the AgeI sequence at the 5′ end and the XbaI sequence at the 3′ end and then ordered for synthesis from Integrated DNA Technologies (IDT). They were then cloned into the pD2i-3′△AgeIWT plasmid. Immediately flanking these restriction sequences, two primer annealing regions were inserted to allow PCR amplification of sufficient template for insertion into pD2i3′△AgeIWT. Wild type and mutant 3′ UTR gBlock sequence design and amplification primers can be found in supplementary data. PCR reactions were cleaned up using Promega’s Wizard DNA cleanup kit according to the manufacturer’s protocol (Promega A7280).

### Viral infections of mammalian and mosquito cells and Northern blot analysis

Vertebrate cells were incubated in trypsin (Corning 25-053-Cl) for ∼5 minutes and mosquito cells were scraped for removal from flask. Suspended cells were washed 2X in PBS and quantified prior to plating in 6-well plates in cell-specific media. Plates were incubated at the appropriate temperature for 3-4 hours to allow re-adherence to plate surface. P1 virus was thawed and added to cell supernatant for a multiplicity of infection (MOI) of 1. Infections were incubated at appropriate temperature or 1 hour with mixing/swirling every 15 minutes. At 1 hour, infection supernatant was removed, replaced with 2 mL cell-specific media and plates incubated for the indicated time. For harvesting of RNA for Northern analysis, cells were lysed in RLT buffer (Qiagen) and processed according to the RNA cleanup protocol of the Qiagen RNeasy kit. For Northern blot analysis sample preparation, 2 ug total RNA was diluted in RNase-free H2O to 13 μL total volume, mixed 1:1 with 2X RNA loading dye, denatured for 5 minutes at 80°C and then placed on ice prior to loading. 6% TBE-Urea gels (Invitrogen EC6865BOX) were pre-run for 30 minutes at 160 Volts (V) prior to sample loading. Samples were run for 20 minutes at 30 V, then 50 minutes at 160 V. Gels were stained with ethidium bromide and imaged before RNA transfer to a positively charged nylon membrane (Fisher Scientific RPN203B) and UV crosslinking. Membranes were then transferred to glass hybridization tubes with 10 mL Ultrahyb-oligo hybridization buffer (Invitrogen AM8663) and rotated for 1 hour at 42°C. Radiolabeled probe in hybridization buffer was exchanged onto the membrane and incubated for a minimum of 14 and maximum of 24 hours at 42°C. Following incubation, membranes were washed for 10 minutes 4x in 2X SSC with 1% SDS (30 mM NaCl, 3 mM sodium citrate, pH 7) and then placed in an exposure cassette with a phosphor screen for imaging. 1 μmol DNA oligo probes complementary to the dumbbell region (Sequence: 5′-CCGCTAGTCCACTACGCCATGCGTACAG-3′), U6 snRNA (Sequence: 5′-TATGGAACGCTTCACGAATTTGCGTCATCC-3′), or Aedes 5.8S rRNA (Sequence: 5′- ATGCGTTCAACGTGTCGG-3′) were radiolabeled via T4 polynucleotide kinase (PNK, NEB M0201L) reaction with 2 μL 5 mCi [γ^32^P] ATP (PerkinElmer).

### Sequence alignment and covariation model

An initial seed alignment of tandem xrRNAs was manually created by obtaining sequences of representative class 1a xrRNAs from ZIKV (NC_012532), WNV1 (NC_009942), WNV2 (NC_001563) and DENV-1 (NC_001474) spanning from the 5′ end of xrRNA1 through the 3′ end of xrRNA2. Each of the two xrRNA regions was aligned individually based on previous structural and bioinformatic data (44). This preliminary seed alignment was used to query a RefSeq viral database (downloaded from NCBI on 01/22/2019) using Infernal v1.1 (79). A total of 17 sequences had two sequential intact and full-length xrRNAs and were retained in this curated tandem xrRNA alignment (see Supplemental Material). The region between xrRNA1 and xrRNA2 is known to contain a separate structured element (stem loop III; SLIII) in some viruses and the base pairing for this element was aligned based on previous experimental data (80). The resulting alignment was used to generate a consensus model calculated using R-Scape (81), graphically depicted in R2R (82), and adjusted in Adobe Illustrator.

### CryoEM imaging and data processing

Sample preparation and data collection: CryoEM grids of DENV2 tandem xrRNA samples were prepared as previously done for RNA-only constructs of similar size (83). Briefly, *in vitro*-transcribed RNA was folded at 18 µM in 1 mM MgCl2 and 50 mM Na-MOPS pH 7 by incubating at 80°C for 1.5 minutes, followed immediately by incubating in ice for several minutes. After this, MgCl2 solution (90 mM) was added to bring the sample to the final imaging [Mg^2+^] of 10 mM. C-flat holey carbon grids (1.2 µm hole diameter; 1.3 µm spacing; 400 mesh copper grid) were cleaned using a Gatan Solarus Model 950 advanced plasma system with the following settings: ‘Cleaning time’: 6 seconds, ‘Vacuum target’: 70 mTorr, ‘Vacuum range’: 0 mTorr, ‘Pumping switch point’: 20 Torr, ‘Turbo pump speed’: 750 Hz, ‘O2 gas flow’: 27.5 sccm, ‘H2 gas flow’: 6.4 sccm, ‘Air gas flow’: 0.0 sccm, ‘Gas flow timeout’: 20 seconds, ‘Forward RF target’: 50 W, ‘Forward 7 RF range’: 5 W, ‘Maximum reflected RF’: 5 W, ‘RF tuning timeout’: 4 seconds, ‘RF tunning attempts’: 3. A 3 µL drop of folded RNA was deposited on plasma-cleaned grids and a FEI Vitrobot Mark IV (temperature: 4°C; humidity: 100%) was used to freeze the grids using liquid ethane, using a blot time of 0.5 seconds and a blot force of -5. Imaging was done on a 200 kV ThermoFisher Talos Arctica equipped with a Gatan K3 Summit direct electron detector, using a pixel size of 0.89 Å and a total dose of 113.9 electrons/Å2. Leginon software was used to collect 1490 movies.

Data processing: CryoSPARC was used for data processing (**Fig. S5**). Motion-corrected micrographs were generated using ‘Patch Motion Correction’ and CTF parameters were obtained using ‘Patch CTF Estimation’. To generate initial templates for particle picking, we manually picked 3,817 particles, extracted them using a box size of 288 pixels, and used 2D classification to generate twenty-five 2D templates. These templates were then imported into a ‘Template Picker’ job to pick particles in a subset of 200 micrographs. This resulted in 25,155 particles (box size: 288 pixels) that were classified into twenty-five 2D classes that were used as templates to pick particles in the full dataset of 1,490 micrographs. After ‘Inspect Particle Picks’ jobs, particle extraction (box size: 288 pixels) and 2D classification, this resulted in 166,959 particles that we used in an ‘*Ab Initio* 3D Reconstruction’ job to generate three volumes. Only one of the three reconstructions resulted in a volume consistent with grooves resembling RNA secondary structural features, the expected size of the construct, and a pseudo-2D symmetry consistent with tandem xrRNAs. In addition, this volume was consistently generated by different ‘*Ab Initio* 3D Reconstruction’ jobs with a variable number of classes (**Fig. S5**) These observations suggested that this volume represented the major conformational state of the DENV2 tandem xrRNA construct. We then used a series of ‘Heterogenous Refinements’ to classify the particles and refine this volume, as done previously (83). This resulted in 26,585 particles that we used in a ‘Homogenous Refinement’ job to generate the final reconstruction with an overall resolution of 7.0 Å.

### Nonsense-mediated RNA decay (NMD)-based reporter

The NMD surrogate reporter plasmids were adapted from pcDNA5/FRT/TO-globin_del5UTR_WT-xrRNA-4H or pcDNA5/FRT/TO-globin_del5UTR_PTC39-xrRNA-4H (63) for mammalian or pAc5.1b (68) for insect transfections respectively. Each DENV2 3ʹ UTR construct was designed as a double-stranded DNA gBlock (IDT) and was subsequently amplified by PCR using custom DNA primers and recombinant Phusion Hot Start polymerase (New England Biolabs). The amplified product was confirmed by gel electrophoresis to be the correct size and cloned into the XhoI and NotI sites downstream of the β-globin gene in the mammalian system or cloned by Genscript downstream of the alcohol dehydrogenase gene for the insect system. Cloned plasmids were amplified in competent *E. coli* DH5α cells and purified via the use of a Qiagen miniprep kit (Qiagen). The recovered plasmid stocks were verified through sequencing (Eton Biosciences or Quintara Biosciences).

### Mammalian cell transfections

2 mL of Vero cells were plated in 6 well plates at a cell count of 2.5×10^5^ cells/mL. Cells were monitored 24 to 48 hours post plating for growth until ∼90% confluency. The cells were then transfected with 2.5 ug of plasmid using the Lipofectamine 3000 kit (ThermoFisher Scientific). At 48 hours post transfection, whole RNA was extracted from the cells using the Qiagen RNeasy Mini Kit (Qiagen).

### Insect cell transfections

C6/36 cells were plated in T-25 flasks at a cell count of 7×10^5^ cells/mL. Cells were monitored for up to 72 hours post plating for growth and media changes until ∼90% confluency. The cells were then transfected with 1 ug of plasmid using the Lipofectamine 3000 kit (ThermoFisher Scientific). At 48 hours post transfection, whole RNA was extracted from cells using the Qiagen RNeasy Mini Kit (Qiagen).

### Northern analysis of reporter assays

1.2 µg of total RNA from cells was treated with RNaseH by using the DNA/RNA hybrid oligo that targeted the 3ʹ end of the DENV 3ʹ UTR (5ʹ-mCmCmAmGmCmGmUmCmAmATATGmCmUmGmUmUmUmUmUmUmG-3 ʹ) to remove the polyA tail in a 10 μL total volume reaction and then mixed 1:1 with 2x RNA loading buffer and run on a 6% denaturing PAGE gel (Invitrogen) with RiboRuler RNA ladder (Thermo Scientific). Gels were pre-run for 30 minutes at 160 V prior to sample loading. Samples were run for 20 minutes at 30 V, then 50 minutes at 160 V. Gels were then stained with ethidium bromide and imaged. The RNA from the gels was subsequently transferred to a HyBond-N+ nylon membrane (GE Life Sciences) using an electrophoretic transfer apparatus (Idea Scientific). The membrane was crosslinked using a UV Stratalinker and transferred to glass hybridization tubes and blocked at 42°C using ULTRAhyb Oligo hybridization buffer (Invitrogen AM8663) for 20 minutes while rotating. Radiolabeled probe in hybridization buffer was exchanged onto the membrane and incubated for a minimum of 18 and maximum of 24 hours at 42°C. Following incubation, membranes were washed for 10 minutes 4x in 2X SSC with 1% SDS (30 mM NaCl, 3 mM sodium citrate, pH 7) and then placed in an exposure cassette with a phosphor screen for imaging. DNA oligo probes complementary to the dumbbell region (Sequence: 5′-CCGCTAGTCCACTACGCCATGCGTACAG-3′) or U6 RNA (Sequence: 5′-TATGGAACGCTTCACGAATTTGCGTCATCC-3′ were radiolabeled via T4 polynucleotide kinase (PNK, NEB M0201L) reaction with 2 μL 5 mCi [γ^32^P] ATP (PerkinElmer).

### Generation of RNAs for chemical probing

Each DENV2 3ʹ UTR construct was designed as a double-stranded DNA gBlock (IDT) and was subsequently amplified by PCR using custom DNA primers and recombinant Phusion Hot Start polymerase (New England Biolabs). The amplified product was confirmed by gel electrophoresis to be the correct size and subsequently purified using the PCR cleanup kit (Promega). The amplified DNA was then *in vitro* transcribed using the NEB Highscribe kit for 3 hours at 37°C. 2 μL of DNase1 (NEB) was added and the transcription reactions were incubated at 37°C for an additional 15 minutes. RNA was then cleaned up using the Monarch kit (NEB) following the standard protocol with a few minor changes. First, 30 μL of DEPC water was added to the transcription reactions before beginning cleanup. After the second wash step (step 5) another spin was done with the lid off to completely remove EtOH from the columns. 30 μL of DEPC water was added to the columns and they sat on the benchtop for 5 minutes before spinning/eluting as this drastically increased the yields.

### Chemical probing

Chemical probing of the *in vitro*-transcribed mutant DENV2 3ʹUTRs was done by adapting the previously described protocol (70). Briefly, 2 µg of RNA was re-folded by heating to 95°C for 1 minute then cooled to room temperature. After cooling, 6 μL of 3.3x folding buffer (333 mM HEPES (pH 8), 333 NaCl, 33 mM MgCl2) was added to the RNA and incubated at 37°C for 20 minutes. Following incubation the RNA was split in half and 1 µg of RNA was modified with either 10 mM 1-methyl-7-nitroisatoic anhydride (Millipore Sigma) or DMSO for 75 seconds at 37°C. Modified and control samples were purified and concentrated using the RNeasy Micro Kit (Qiagen). Following modification and purification, RNA was incubated with 200 ng/µL Random Primer 9 (NEB) and 1 µL of a reverse primer that was specific to the DENV2 3′UTR at 65°C for 5 minutes and then cooled on ice. 8 µL of MaP buffer (125mM Tris pH 8.0, 187.5 mM KCl, 15 mM MnCl2, 25 mM DTT), 1.23 mM equimolar dNTP mix, and 200 U Superscript II was added to each reaction and incubated in a thermocycler at 25°C for 2 minutes, 25°C for 10 minutes, 42°C for 3 hours, 70°C for 15 minutes, and cooled to 4°C. The RNA/cDNA hybrids were then purified on G-25 microspin columns (Cytiva) following the manufacturer’s protocol with slight modification. To ensure there was no contamination of carryover DNA to the columns they were extensively washed with 500 μL of deionized water (3 washes each). Second-strand synthesis was carried out using the NEBNext Ultra II Non-Directional RNA Second Strand Synthesis Module (NEB) following the standard protocol. The resultant cDNA was purified using the DNA Clean & Concentrator-5 kit (Zymo). The resultant dsDNA pool was quantified with a Qubit 3 Fluorometer (Thermo) using the Qubit dsDNA HS Assay Kit (Thermo). Next-generation sequencing library prep was performed with the Nextera XT DNA library preparation kit (Illumina) using the Nextera XT Index Kit v2 Set B (Illumina). Samples were normalized and pooled together using a combination of the Qubit and the 4200 TapeStation System (Agilent Technologies) with the High Sensitivity D5000 Screen Tape (Agilent Technologies). Samples were sent to Novogene for sequencing on the NovaSeq6000 asking for 10 million paired end reads for each sample. For SHAPE-Map analysis, raw sequencing reads were first visualized with FastQC (v0.11.9) to assess the quality of the data. The raw reads were then trimmed using Trimmomatic (v040) to remove any residual Illumina adapters. The processed reads were then analyzed with Shapemapper2 with the default parameters and a minimum read depth of 5000 reads. The resultant data was analyzed using a custom in-house pipeline of Python (v3.9.10) and R (v4.2.2) scripts. Average reactivities of two or three biological replicates per construct were then plotted onto the proposed secondary structure using RNAcanvas (v1.1.16) (84).

### Generation of RNAs for SAXS

Each DENV2 tandem xrRNA construct was designed as a double-stranded DNA gBlock (IDT) and was subsequently amplified by PCR using custom DNA primers and recombinant Phusion Hot Start polymerase (New England Biolabs). DNA was subsequently *in vitro* transcribed in 5 mL reactions with an in-house prepped RNase inhibitor (expressed and purified from the pMAL-c5X-MBP-RNase Inhibitor plasmid (#153314) from Addgene). RNA was spun down to remove free phosphates and supernatant was transferred into a new tube with 3x volume chilled EtOH. RNA was then pelleted and resuspended in 2.5 mL 2X loading buffer and run on an 8% polyacrylamide gel. Bands were excised and eluted in 50 mL of 20 mM sodium acetate pH 5.2 and rocked at 4°C overnight. The elutions were then concentrated in 10,000 MWCO Amicon spin filters (Sigma) and quantified via Nanodrop.

### SAXS collection and analysis

SEC-SAXS measurements (I(q) vs. q, where q is defined in the equation*q* = 4*πsinθ*/*λ* and 2θ is the scattering angle and λ the 31 X-ray wavelength) were performed at the 16-ID (LiX) beamline at the National Synchrotron Light Source II (NSLS-II) equipped with a Pilatus 1M detector, using the beamline’s standard SAXS configuration as described by Yang et al. (85). RNAs were re-folded by heating to 95°C for 1 minute, then 2 or 4 mM MgCl2 was added (for samples with Mg) and then cooled at room temperature for 5 minutes. Final RNA concentration was 2 mg/mL. 85 µL of the RNA sample was loaded onto a Superdex 200 Increase 5/150 GL (GE Healthcare) column pre-equilibrated in SAXS buffer with or without MgCl2 using an Agilent HPLC 4 system at a flow rate of 0.75 mL/minute. 600 successive 2D SAXS data frames were collected from the continuously flowing eluate. Data averaging, processing and visualization were done with BIOXTAS RAW suite and the ATSAS 4.0.0-2 software suite. We used CHROMIXS for the selection of sample and buffer frames and PRIMUS for calculation of all reported parameters and generation of Guinier, pairwise distance distribution function, and Kratky plots.

### Human Xrn1 protein expression and purification

The gene encoding for the N-terminus (residues 1-1180) of Human Xrn1 (GenBank: AAN11306.1) with a C-terminal His6 tag was synthesized and cloned into a pET28 bacterial expression vector (Genscript). For expression, the cloned plasmid was transformed into *E. coli* strain Rosetta (DE3)pLysS and grown in LB medium at 37°C. Cell growth was monitored by absorbance at 600 nm, cells were cooled and expression was induced upon reaching 0.6 at OD600 by adding IPTG to a final concentration of 0.5 mM. Temperature was lowered to 18°C and cells were harvested ∼16 hours later. Harvested cells were pelleted and frozen at -20°C for storage. For purification, frozen cell pellets were thawed and resuspended in lysis buffer (500 mM NaCl, 10 mM imidazole, 50 mM HEPES [pH 7.4], 2.5 mM βME, 10% glycerol, 5 mM MgCl2, 5 mM ATP, and 1X EDTA-free protease inhibitor [Roche]). Resuspended cells were lysed by sonication and the resulting lysate was clarified by centrifugation (20,000 x g for 30 minutes). Clarified lysate was applied to a Ni affinity column (Pierce High Capacity Ni-IMAC Resin, EDTA-compatible) that was equilibrated with lysis buffer (sans protease inhibitor). Bound protein was washed with 20 column volumes of lysis buffer (sans protease inhibitor and ATP / MgCl2), 20 column volumes of high salt buffer (1 M NaCl, 10 mM imidazole, 50 mM HEPES [pH 7.4], 2.5 mM βME, 10% glycerol), 20 column volumes of lysis buffer (sans protease inhibitor), and 20 column volumes of low salt buffer (100 mM NaCl, 50 mM HEPES [pH 7.4], 2.5 mM βME, 10% glycerol). Protein was eluted from Ni-IMAC resin with a buffer containing 100 mM NaCl, 300 mM imidazole, 50 mM HEPES [pH 7.4], 2.5 mM βME, and 10% glycerol. Eluate was concentrated and loaded onto a HiTrap Q HP anion exchange column that was equilibrated with Anion Exchange Buffer A (100 mM NaCl, 50 mM HEPES [pH 7.4), 2.5 mM βME). After application, the Q column was washed with 10 CV of Anion Exchange Buffer A before being eluted with a linear gradient from 50 mM to 1 M NaCl. Elution was monitored by UV absorbance at 280 nm and fractions were assayed by SDS-PAGE. Fractions with a high 260 nm signal were avoided. Fractions containing pure Xrn1 were pooled, concentrated, and applied to a HiLoad 16/60 Superdex 200 (Cytiva) column that was equilibrated in SD200 buffer (100 mM NaCl, 50 mM HEPES [pH 7.4], 2 mM βME). Elution was monitored by UV absorbance at 280 nm. Peak fractions were assayed by SDS-PAGE, collected, and concentrated to 15-30 µM. Concentrated protein was stored at 4°C for up to 1 month.

### Transcription and purification of DENV2 3ʹUTRs for hXrn1 degradation assays

DNA templates for the DENV2 sfRNA (endogenous nucleotides 10290 through 10723 with an appended 34 nucleotide 5′ leader) and its mutations were ordered as gBlock DNA fragments (IDT). PCR reactions using primers containing an upstream T7 promoter were used to generate dsDNA templates for transcription. PCR conditions were generally as follows: 0.1 ng/µL DNA, 200 µM dNTP, 0.5 µM forward and reverse primers, 1X Phusion reaction buffer, and 0.05 units/µL of Phusion. dsDNA amplification was confirmed by 2% agarose gel electrophoresis. Transcription reactions to generate the sfRNAs were performed using 0.3 µg/µL of dsDNA template, 1X transcription buffer (30 mM Tris-HCl [pH 8], 10 mM DTT, 3.5 mM spermidine, and 0.1% Triton X-100), 7.5 mM rNTPs, 35 mM MgCl2, and T7 RNA polymerase. Reactions were incubated at 37°C for approximately 2 hours (or until cloudy). Insoluble pyrophosphate was removed by centrifugation at 4200 x g for 15 minutes. The RNA-containing supernatant was precipitated with three volumes of -20°C 100% EtOH and incubated at -20°C for 1 hour. The precipitated RNA solution was centrifuged at 4200 x g for 30 minutes at 4°C to pellet the RNA. The EtOH solution was removed and the resulting RNA pellet was allowed to dry. The RNA was resuspended in 8 M urea loading buffer and purified by denaturing 8% Urea-PAGE. Bands were visualized by UV back-shadowing and excised from the Urea-PAGE gel. RNA was removed by the excised bands via the “crush-and-soak” method using RNase-free water containing 300 mM sodium acetate (pH 5.4). The RNA-containing solution was filtered using 0.2 micron filters, concentrated using spin concentrators (Amicon), and buffer exchanged into RNase-free water using Zeba desalting spin columns. RNA concentrations were determined using absorbance at 260 nm and RNA was stored at -20°C until use.

### *In vitro* exoribonuclease resistance assays

Xrn1 degradation assays followed previously described protocols (42). Prior to any degradation assay, sfRNAs (5 µM in 36 µL) were refolded in 100 mM NaCl, 2.5 mM MgCl2, and 10 mM HEPES (pH 7.4) using a thermocycler programmed to heat to 90°C for 2 minutes, 20°C for 5 minutes, and cooled to 4°C until use. DTT was added to RNA samples at a final concentration of 1 mM. Following refolding, 2 µL of 0.8 mg/mL BdRppH (42) was added to each sample, which was split between two tubes. In one tube, 1 µL of buffer (100 mM NaCl, 2.5 mM MgCl2, and 10 mM HEPES [pH 7.4]) was added to serve as a (-) Xrn1 control. In the other tube, 1 µL of 1 mg/mL Xrn1 was added. All samples were incubated at 37°C for 2 hours. Reactions were quenched by the addition of an equal volume of Urea loading buffer (8 M Urea, 30 mM EDTA, xylene cyanol and bromophenol blue). Samples were heated at 98°C for 5 minutes prior to loading on a 6% Urea-PAGE gel for analysis. Gels were run at 180 V for 90 minutes, stained with 0.1% methylene blue for 3 minutes, destained in water, and imaged on a Bio-Rad Gel Doc EZ Imager. Assays were performed three times to confirm the Xrn1-dependent banding patterns.

## ACKNOWLEDGMENTS

The authors thank K. Segar, S. Zangari, and A. MacFadden for assistance with preparation of some reagents, D. Costantino for technical support and manuscript proofreading, and all current and former members of the Kieft Lab for useful discussions and input. Thank you to Drs. L. van Dyk and E. Clambey for input, support, and the use of laboratory space. This research used the LiX beamline (16-ID) of the National Synchrotron Light Source II, a U.S. Department of Energy (DOE) Office of Science User Facility operated for the DOE by Brookhaven National Laboratory under Contract No. DE-SC0012704. The LiX beamline is part of the Center for BioMolecular Structure (CBMS), which is primarily supported by the National Institutes of Health, National Institute of General Medical Sciences (NIGMS), through a Center Core Grant (P30GM133893), and by the DOE Office of Biological and Environmental Research (KP1607011). LiX also received additional support from NIH Grant S10OD012331. Dr. S. Chodankar at beamline 16-ID (LiX) assisted with SAXS data collection. We thank Drs. L. Pollack and S. Korn for advice regarding SAXS data analysis. CryoEM data was collected at the University of Colorado Anschutz Medical Campus facility, supported in part by grant P30CA046934 and then managed by Dr. E.R Camacho. This work is supported by NIH grants F31AI176728 (E.S.), T32AI052066 (E.S. and Z.O.), F32GM139385 (S.L.B.) and R35GM118070 (J.S.K.). M.E.S. was supported by a Jane Coffin Childs postdoctoral fellowship and S.L.B. was an HHMI Hanna Gray Fellow.

