## Supplemental Figures for "An A-rich linker between dengue virus tandem xrRNAs facilitates functional coordination"

### Supplemental figures contents:

**Supplemental Figure S1:** Diagrams of mut-xrRNA1, mut-xrRNA2, swap #1, swap #2, and +32 spacer mutants

**Supplemental Figure S2:** Nonsense-mediated decay (NMD) reporter systems design and protocol

**Supplemental Figure S3:** Sequence alignment of tandem xrRNAs

**Supplemental Figure S4**: SHAPE-MAP analysis of wild type and swap #1 mutant tandem xrRNAs

**Supplemental Figure S5:** Overview of the data processing workflow for single-particle cryoEM analysis of the DENV2 tandem xrRNA

**Supplemental Figure S6:** Small-angle x-ray scattering analysis plots of wild type and swap #1 mutant tandem xrRNAs

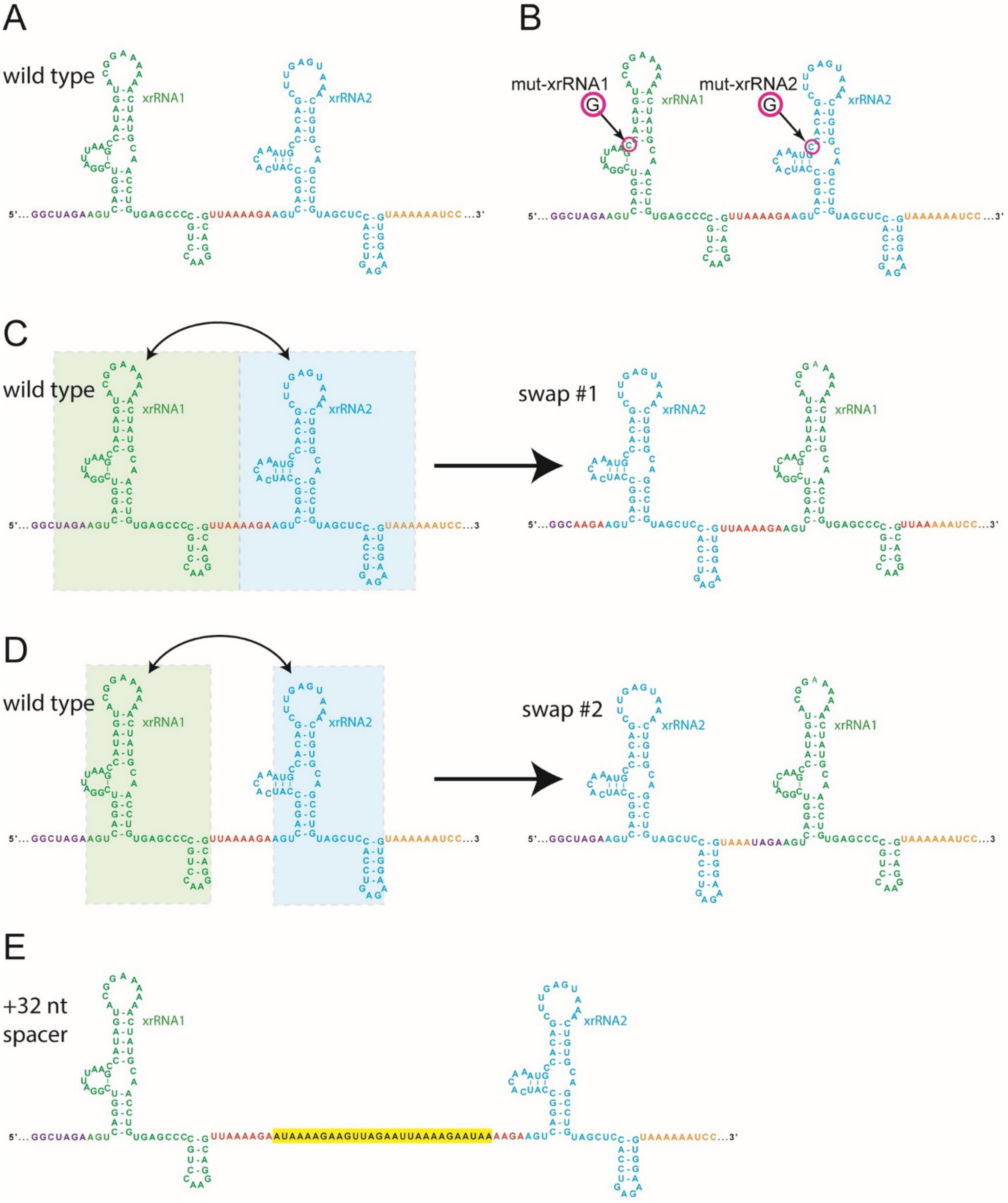


**Supplemental Figure S1. Diagrams of mut-xrRNA1, mut-xrRNA2, swap #1, swap #2, and +32 spacer mutants.** (A) Secondary structure diagram of wild type DENV2 tandem xrRNAs. Colors indicate various elements in the RNA. (B) Location of point mutations resulting in mut-xrRNA1 and mut-xrRNA2. Mut-xrRNA1+2 contains both point mutations. (C) Left: wild-type tandem xrRNA with the parts that were swapped to make swap #1. Right: swap #1 mutant, with sequences colored to show what was swapped. (D) Same as panel C, but for swap #2 RNA. (E) Diagram of the +32 nt spacer mutant with the added sequence highlighted yellow.

Reporter design and protocol:

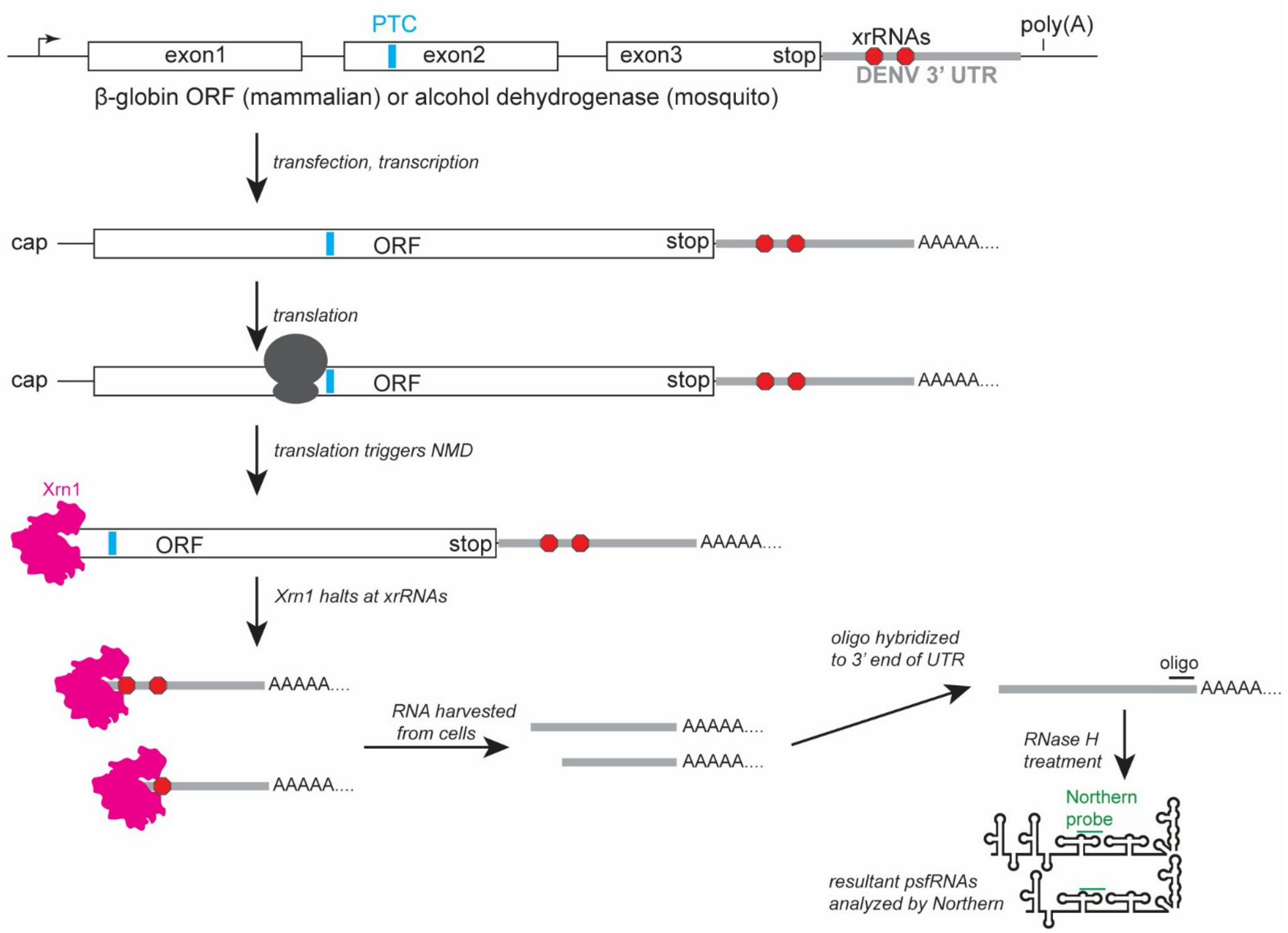


**Supplemental Figure S2. Nonsense-mediated decay (NMD)-based reporter systems design and protocol.** The system comprised expression plasmids containing either the β-globin gene (mammalian system) or the alcohol dehydrogenase gene (mosquito system), with a premature termination codon (PTC, cyan) located within the second exon. Wild type or mutant tandem xrRNAs were placed downstream of these genes. Transfection of appropriate cell types with the plasmid led to transcription and processing to yield mature mRNA. Translation triggered NMD induced by the PTC, leading to recruitment and degradation by Xrn1 (magenta). Xrn1 contact with the xrRNA then led to accumulation of pseudo-sfRNAs (psfRNAs, so named as they are not produced by viral infection). Total RNA was then harvested 48 hours post infection, and a DNA oligonucleotide complementary to the 3' end of the DENV2 3' UTR was used to target RNase H to remove the poly(A) tail; this was necessary as the poly(A) tail was heterogeneous, leading to smeared bands on the gel. The resultant RNA was then analyzed by Northern blot, using a probe complementary to sequence in the first dumbbell structures downstream of xrRNA2 (green).

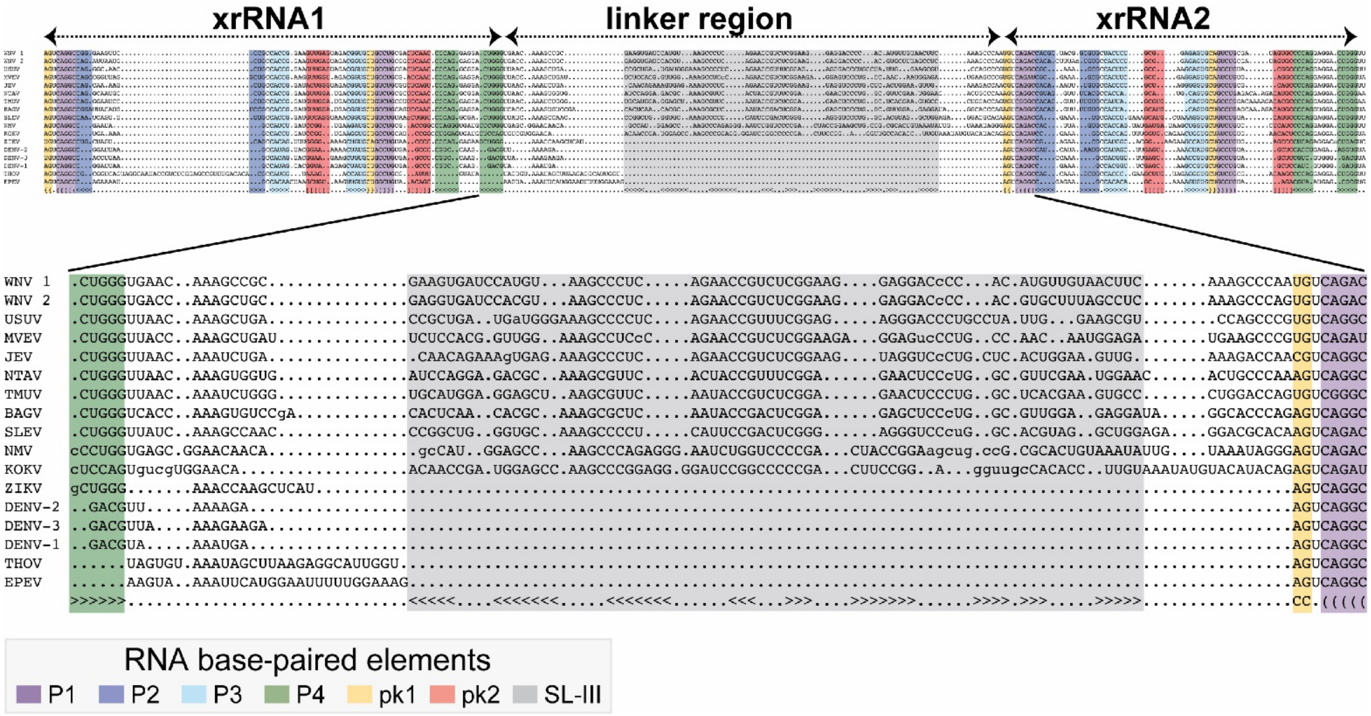


**Supplemental Figure S3. Sequence alignment of tandem xrRNAs.** Alignment based on secondary structure and sequence of all orthoflaviviruses with a tandem arrangement of xrRNAs. Nucleotides were colored to depict base pairing of conserved secondary structure features.

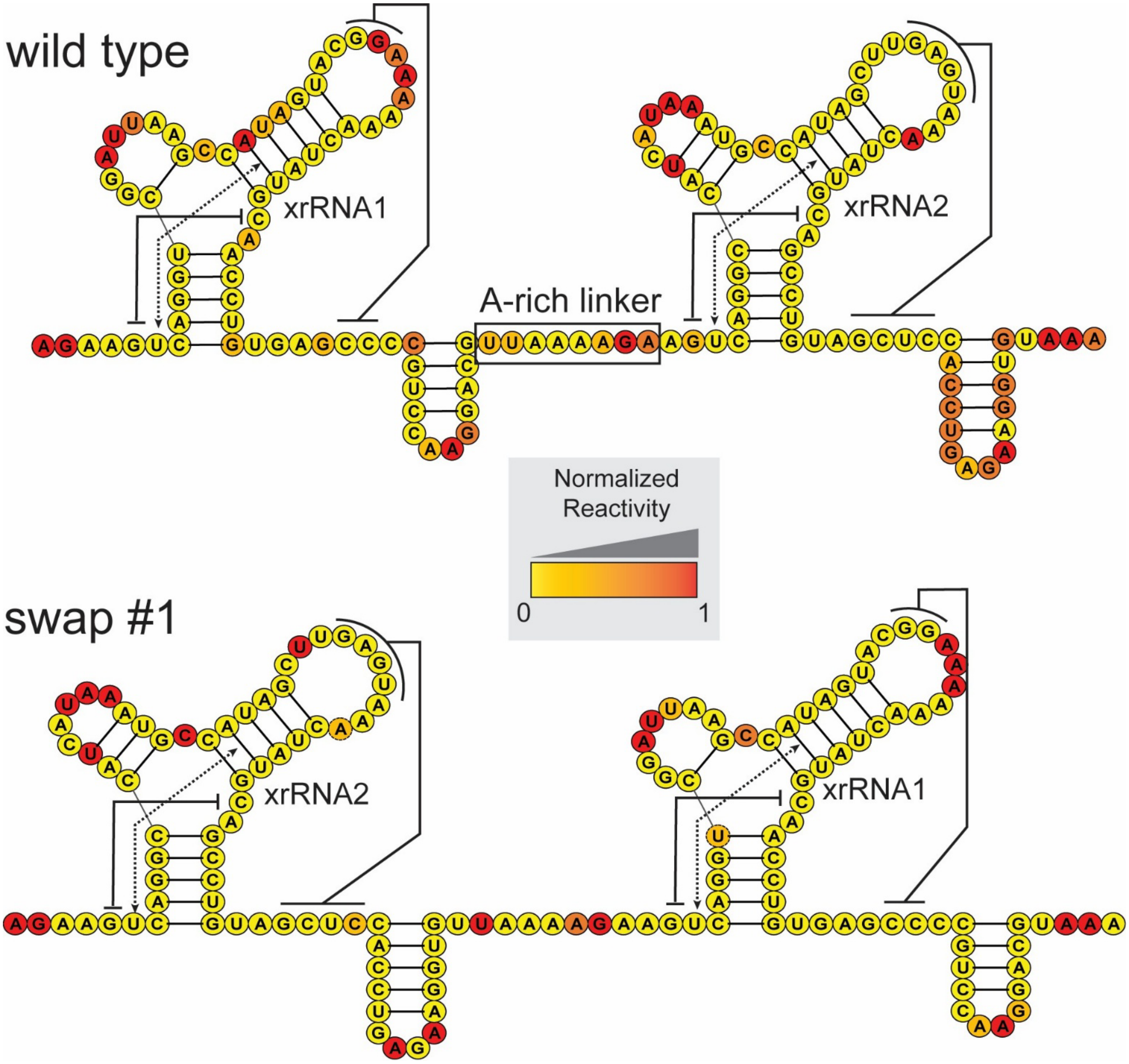


**Supplemental Figure S4. SHAPE-MAP analysis of wild type and swap #1 mutant tandem xrRNAs.** Secondary structure diagram of the WT (top) and swap #1 (bottom) DENV2 tandem xrRNAs in the context of the full 3′ UTR with the results of 1M7 chemical probing. Key tertiary contacts are diagrammed with Watson-Crick base pairing shown with solid lines and a base triple shown with a dashed line.

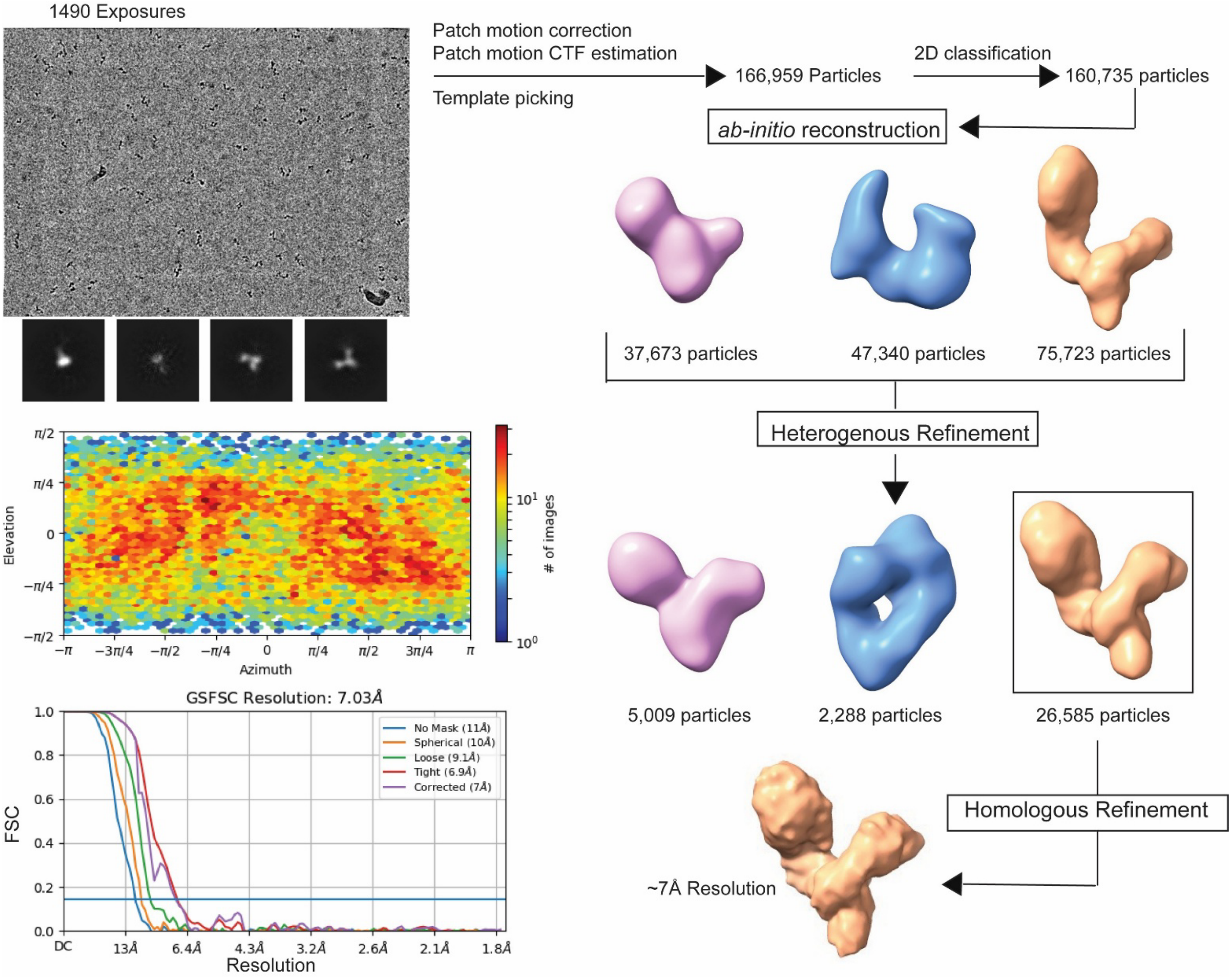


**Supplemental Figure S5: Overview of the data processing workflow for single-particle cryoEM analysis of the DENV2 tandem xrRNA.** The cryoEM processing pipeline in cryoSPARC for the DENV2 tandem xrRNA using data collected with a 200 kV electron microscope. Using cryoSPARC, particles were picked and put through 2D classification to remove junk particles and any low-resolution 2D classes. *Ab initio* reconstruction and heterogenous refinement further removed junk particles and allowed for model building of the tandem xrRNAs. The FSC graph for local and global resolution is shown for the final model used for model building.

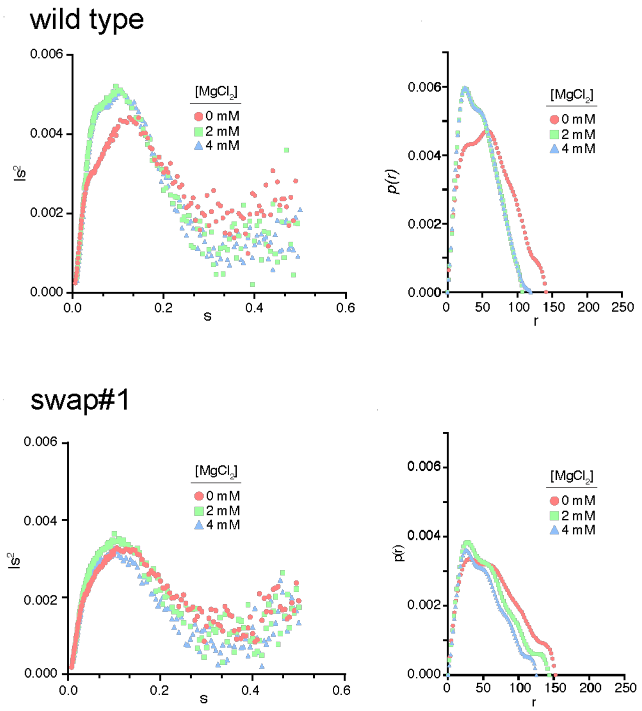


**Supplemental Figure S6. Small-angle x-ray scattering analysis plots of wild type and swap #1 mutant tandem xrRNAs.** Small-angle x-ray scattering analysis plots of wild type and swap #1 mutant tandem xrRNAs. Kratky plot (left) or distance distribution function plot (right) for either the WT or the swap #1 tandem xrRNA constructs, comparing data for 0, 2, or 4 mM $Mg^{2+}$.
